# Role for the Na^+^/K^+^ ATPase pump alpha 3 (*ATP1A3*) subunit in folding and lamination of the human neocortex

**DOI:** 10.1101/2020.10.03.319137

**Authors:** Richard S. Smith, Marta Florio, Shyam K. Akula, Jennifer E. Neil, Yidi Wang, R. Sean Hill, Melissa Goldman, Christopher D. Mullally, Nora Reed, Luis Bello-Espinosa, Laura Flores-Sarnat, Fabiola Paoli Monteiro, Casella B. Erasmo, Filippo Pinto e Vairo, Eva Morava, A. James Barkovich, Joseph Gonzalez-Heydrich, Catherine A. Brownstein, Steven A. McCarroll, Christopher A. Walsh

## Abstract

Osmotic equilibrium and membrane potential in animal cells depend on concentration gradients of sodium (Na^+^) and potassium (K^+^) ions across the plasma membrane, a function that is catalyzed by the Na,K-ATPase alpha subunit. In vertebrates, four paralogous genes, *ATP1A1-4*, encode distinct alpha subunit isoforms (*a_1_-a_4_*), three of which (*a_1_, a_2_, a_3_*) are expressed in the brain, and two (*a_1_, a_3_*) predominantly in neurons. The *a_3_* isoform, encoded by *ATP1A3*, is critical to neuronal physiology, and a growing spectrum of neurological diseases are associated with *ATP1A3* pathogenic variants, with ages of onset ranging from early childhood to adulthood. Here, we describe *ATP1A3* variants encoding dysfunctional *a_3_* subunits in children affected by polymicrogyria, a developmental malformation of the cerebral cortex characterized by abnormal folding and laminar organization. To gain cell-biological insights into the spatiotemporal dynamics of prenatal *ATP1A3* expression, we established a transcriptional atlas of *ATP1A3* expression during cortical development using mRNA *in situ* hybridization and transcriptomic profiling of ~125,000 individual cells with single-cell RNA sequencing (Drop-Seq) from various areas of the midgestational human neocortex. We find that fetal expression of *ATP1A3* is restricted to a subset of excitatory neurons carrying transcriptional signatures of neuronal activity and maturation characteristic of the developing subplate. Furthermore, by performing Drop-Seq on ~52,000 nuclei from four different areas of an infant human neocortex, we show that *ATP1A3* expression persists throughout early postnatal development, not only within excitatory neurons across all cortical layers, but also and more predominantly in inhibitory neurons, with specific enrichment in fast-spiking basket cells. In addition, we show that *ATP1A3* expression, both in fetal and postnatal neurons, tends to be higher in frontal cortical areas than in occipital areas, in a pattern consistent with the rostro-caudal maturation gradient of the human neocortex. Finally, we discover distinct co-expression patterns linking catalytic α subunit isoforms (*ATP1A1,2,3*) and auxiliary isoforms (*ATP1B1,2,3*), suggesting the ATPase pump may form non-redundant, cell-type specific α-β combinations. Together, the importance of the developmental phenotypes and dynamic expression patterns of *ATP1A3* point to a key role for *a3* in the development and function of human cortex.

## INTRODUCTION

Generating a folded, 6-layered cerebral cortex relies on the integration of signals from a variety of cell types, coordinating schedules of cell proliferation, migration, differentiation, and maturation. These processes require proper maintenance of ionic gradients and cell membrane potential (Vm) and thus rely on the precise function of ion channels and pumps. Pathogenic mutations in genes encoding ion channels can cause malformations of cortical development (MCDs), collectively known as developmental channelopathies(Smith and Walsh, 2020). For example, pathogenic variants in prenatally expressed genes encoding glutamate (*GRIN2B, GRIN1*) and sodium channels (*SCN3A*) can lead to MCDs(Fry et al., 2018; Platzer et al., 2017; Smith et al., 2018), at least in part via gain-of-function, cell-autonomous effects in neurons and potentially other cell types(Smith et al., 2018). These developmental channelopathy variants in glutamate and sodium channels cause pathogenic increases in cationic flux into cells(Smith and Walsh, 2020), affecting Vm in neural progenitor cells and neurons, which has been shown to be sufficient to disrupt neurogenic and cortical lamination processes in experimental models(Hurni et al., 2017; Vitali et al., 2018).

Neuronal physiology, homeostasis, and signaling depend on the electrogenic activity of the Na,K-ATPase, an ionic pump powered by ATP hydrolysis which maintains Na^+^ and K^+^ gradients across the plasma membrane. The Na,K-ATPase complex contains a large catalytic subunit (α) and two auxiliary subunits (β, FXYD)(Holm et al., 2016). In vertebrates, four paralogous genes (*ATP1A1-4*) encode *a* subunits, of which the *ATP1A3* isoform is predominantly expressed in the brain and is critical to the restitution of electrochemical gradients following neuronal excitation, among several other physiological functions(Kim et al., 2004; McGrail et al., 1991; Picton et al., 2017). Pathogenic variants in *ATP1A3* have been increasingly associated with a spectrum of neurological diseases, with phenotypes ranging broadly in age of onset, from the immediate postnatal period throughout adulthood(Brashear et al., 1993; Smedemark-Margulies et al., 2016), suggesting different vulnerabilities to *ATP1A3* dysfunction during childhood brain maturation. These diseases include (in approximate order of onset in childhood development): early infantile epileptic encephalopathy (EIEE); alternating hemiplegia of childhood (ACH); cerebellar ataxia, areflexia, pes cavus, optic atrophy, and sensorineural hearing loss (CAPOS); childhood onset schizophrenia (COS); and rapid-onset dystonia-parkinsonism (RDP)(Brashear et al., 1993; Smedemark-Margulies et al., 2016). Variants in *ATP1A3* associated with these postnatal diseases are generally categorized as heterozygous loss-of-function single nucleotide variants(Arystarkhova et al., 2019; Holm et al., 2016) which, among other deficits, have been shown to lead to depolarized Vm, such as in neurons from AHC patients(Simmons et al., 2018). During development, bioelectric changes can dramatically alter both intrinsic trajectories of individual cells(Levin et al., 2017) and cortical lamination more broadly(Hurni et al., 2017; Vitali et al., 2018), suggesting that bioelectric alterations in early brain development could in part contribute to subsequent *ATP1A3-*related neurological disorders.

Here, we present four individuals with novel *de novo ATP1A3* variants resulting in polymicrogyria (PMG, the most common MCD(Jansen et al., 2016)), epilepsy, and global developmental delay, and consider a previously described individual case report of AHC with polymicrogyria(Gurrieri et al., 2016). The *ATP1A3* variants we observed associated with PMG clustered in transmembrane regions 7 to 8 of the α3 protein. Using mRNA *in situ* analysis and single-cell RNA sequencing (Drop-Seq) in the human fetal cortex at mid-gestation, we show that *ATP1A3* is restricted to a subset of deep-layer neurons located in the subplate. Moreover, using Drop-Seq in the human infant cortex, we show that postnatal expression of *ATP1A3* at this stage of development extends not only to all excitatory neuron subtypes (with cortical layer biases), but mostly predominates within cortical interneurons. The infant cortex also displays a rostral-to-caudal *ATP1A3* gradient, with highest expression levels in the prefrontal cortex. Last, within individual cells, variable levels of ATPase α and β subunit isoforms offers differential vulnerability between excitatory and inhibitory neurons, and may provide a partial explanation for the broad phenotypic range associated with *ATP1A3* mutations.Smith et al.,

## RESULTS

### Individuals with *ATP1A3* variants display cortical malformation phenotypes

Magnetic resonance imaging (MRI) of four unrelated individuals with *de novo ATP1A3* variants revealed a range of PMG severity, from extensive bilateral frontoparietal PMG to unilateral PMG (**Figure 1A)**. Detailed descriptions of each case are included in supplemental text.

**Figure 1.**
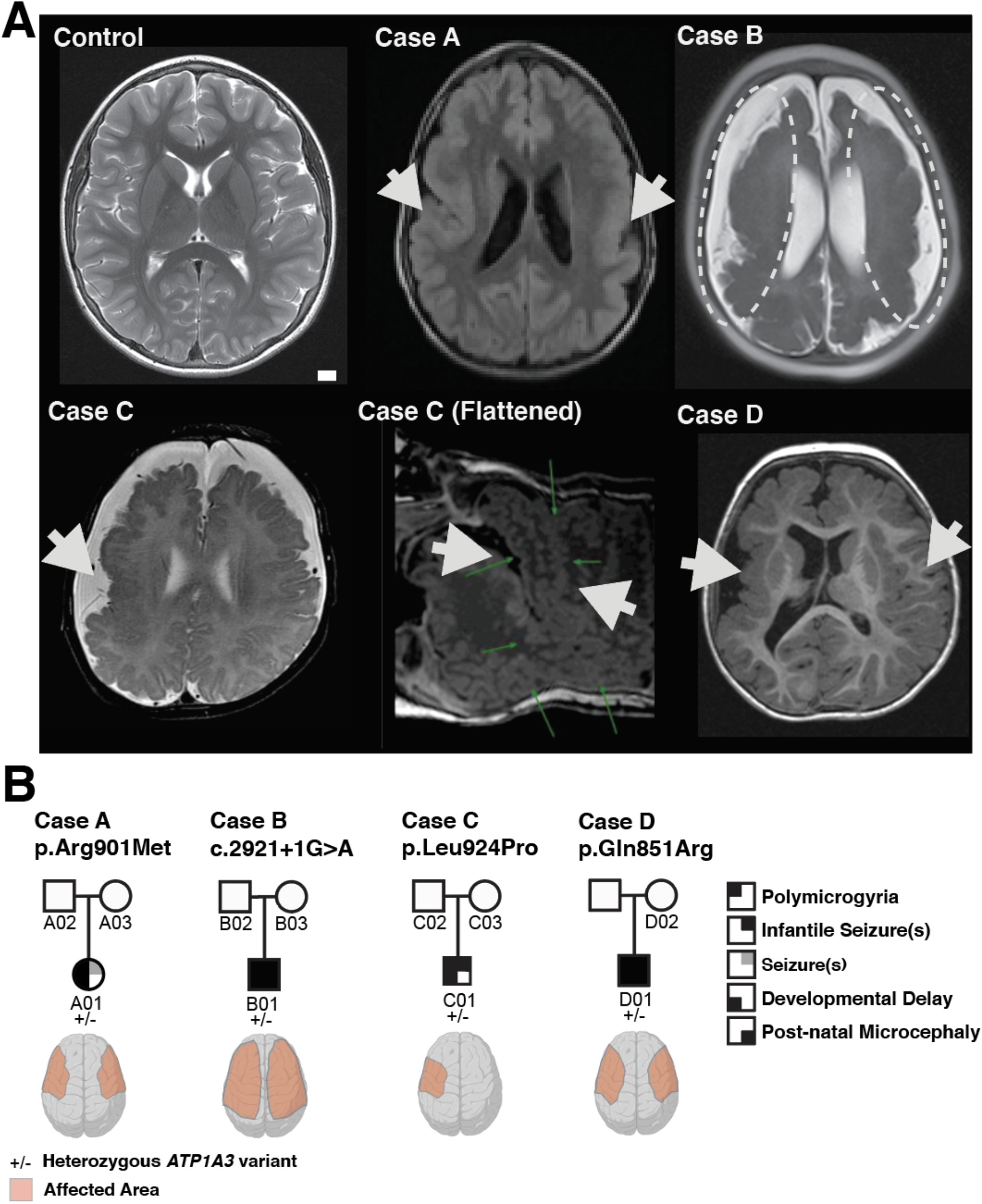
*ATP1A3* variants associated with cerebral cortex malformations. **(A)** MRI images of Case A, B, C and D showing the cortical malformation, polymicrogyria (PMG). White arrows denote gross location of affected brain regions. The Control image comes from an unaffected 11-year-old. Scale bar (1cm) **(B)** Pedigrees with novel *de novo* pathogenic single nucleotide variants in *ATP1A3:* Case A (missense, p.Arg901Met) with bilateral frontoparietal PMG, Case B (splice donor site, c.2921+1G>A) with extensive bilateral PMG, Case C (missense, p.Leu924Pro) with unilateral PMG, and Case D (missense, p.Gly851Arg) with extensive bilateral PMG, more pronounced in the right hemisphere. Individual features: Genotype (−/+, *de novo* heterozygous), PMG, infantile seizures, developmental delay, and postnatal microcephaly shown in pedigree summary. Square, male; circle, female; black and/or gray shading, affected individual. See supplemental text for genetic validation and comprehensive clinical phenotyping.

#### Case A: Bilateral Frontoparietal PMG (p.Arg901Met)

Case A, a female child of nonconsanguineous parents from Portugal, was born at full term and had a short postnatal hospital stay for respiratory distress and jaundice. Brain MRIs performed at 5 months and at 9 years of age revealed extensive bilateral perisylvian polymicrogyria involving the lateral temporal lobes, insulae, and posterior half of frontal and parietal lobes **(Figure 1A**). The posterior part of frontal and temporal cortex appears abnormally thick, with somewhat shallow sulci, giving a pachygyric appearance (bilateral and fairly symmetrical, but with a mild left to right predominance). The most anterior aspects of the frontal and parietal lobes, as well as the medial and inferior cerebral cortex, were spared. At 9 years she developed seizures, manifesting as focal impaired awareness, and was treated with oxcarbazepine. She was developmentally and cognitively delayed, able to sit alone and walk with a walker, point and make some signs but had no speech, and could hold a pencil but not write. Asymmetric spastic quadriparesis (left more significant than right) with hypertonia and athetoid movements of the hands and fingers were noted on examination, as were a high palate, thin upper lip and smooth philtrum. Her head circumference at 9 years was 51 cm (−0.73 standard deviations, SD), and she exhibited drooling and had feeding difficulties. Trio whole exome sequencing (WES) revealed a *de novo ATP1A3* variant, c.2702G>T (p.Arg901Met), absent in both parents (**Figure 1 & S1**).

#### Case B: Extensive Bilateral PMG (c.2921+1G>A)

Case B, a male child of nonconsanguineous parents from the Philippines, was born at 38 weeks’ gestation. His head circumference at birth was 34 cm (−0.83 SD) and he developed seizures 12 hours after delivery. Brain MRI at 6 weeks revealed extensive bilateral polymicrogyria, and small, cyst-like areas were noted within the hippocampal heads and bodies. Video EEGs confirmed a severe epileptic encephalopathy resistant to essentially all antiepileptic drugs. EEGs at 2 months of age showed abundant electroclinical and electrographic seizures beginning predominantly from the right parasagittal region but also from the left and bilateral parasagittal regions. Visual and auditory evoked potential studies were normal at 4 and 5 months of age respectively. Evaluation at 14 months revealed microcephaly, with a head circumference of 41.5cm (−4.63 SD), severe global developmental delay (never rolled over, sat up, or talked), intermittent nystagmus, significant axial and appendicular hypotonia, and hyporeflexia. Trio WES revealed a *de novo ATP1A3* variant, c.2921+1G>A, disrupting a conserved splice site, absent in both parents (**Figure 1A & S1**).

#### Case C: Unilateral PMG (p.Leu924Pro)

Case C, a male child of nonconsanguineous parents from Brazil, was born at 38 weeks’ gestation by Cesarean section for fetal distress. He was hypotonic and noted to have a left clubfoot at birth and was admitted to neonatal intensive care when episodes of upward rolling of the eyes were noted on the first day of life. MRI revealed extensive cortical malformation of the right hemisphere with polymicrogyria comprising the frontal, parietal, and temporal right lobes and the insula, and a small focus of signal abnormality in the periventricular white matter, compatible with recent ischemic injury. EEG revealed the presence of electroclinical seizures and his neurological exam at the time disclosed episodic dystonic posturing of the upper left extremity alternating with excessive movements of closing his left hand with thumb adduction. He was discharged on phenobarbital and levetiracetam. Upon neurological exam at 8 months of age, he had a head circumference of 42.5 cm (−1.75 SD), very poor eye contact, global hypotonia with present reflexes, absent head support, absence of rolling and of object gripping, and presence of babbling sounds. Trio WES revealed a *de novo ATP1A3* variant, c.2771T>C (p.Leu924Pro), absent in both parents (**Figure 1 & S1**).

#### Case D: Extensive multifocal PMG (p. Gln851Arg)

Case D, a male child of nonconsanguineous parents of European descent from the United States, was born at 41 weeks by emergency Cesarean section; the pregnancy was complicated by fetal hydronephrosis. He developed electroclinical seizures and a neonatal encephalopathy requiring cooling and was discharged after 1 month. MRI soon after birth showed multifocal PMG involving both cerebral hemispheres, right greater than left, with the right hemisphere smaller than the left (**Figure 1A)**. At 4 months of age he was admitted for congenital hip dysplasia surgery and noted to have frequent breath holding spells that worsened after the surgery. His varied seizure phenomenology included nystagmus, chin quivering, and hand shaking, which continue intermittently even through phenobarbital, levetiracetam, and oxcarbazepine combination therapy. At 15 and 30 months of age, head circumference was 42.7 cm (−3.5 SD) and 44.2 cm (−3.3 SD), respectively. Physical exam at 18 months demonstrated spastic hemiparesis involving primarily left arm, but including ipsilateral face, and central hypotonia. Other symptoms include feeding difficulties and persistent constipation. WES revealed a *de novo ATP1A3* variant, c.2552A>G (p.Gln851Arg), absent in mother (father was not available for testing).

### PMG-associated *ATP1A3* mutations are predicted to disrupt function

*ATP1A3* encodes the P-type Na,K-ATPase α3 subunit (NM_152296), an integral membrane protein that relies on ATP hydrolysis to transport ions (Na^+^ and K^+^) via 10 membrane-inserted helices (TM1-TM10). This ATPase is functionally supported by the β and FXYD subunits(Holm et al., 2016). *ATP1A3* has several conserved domains across α isoform paralogs (α1-4) and orthologous genes(Clausen et al., 2017), and is a highly constrained gene, with many fewer missense and loss of function variants seen in the general population than predicted (gnomAD Z score of 6.33 and pLI score of 1.0 respectively). Taken together with *ATP1A3’s* known disease association with AHC, RDP, IEE, and CAPOS(Brashear et al., 1993), this suggest the newly identified variants in *ATP1A3* are likely damaging **(Table S1).**

PMG-associated *ATP1A3* variants (p.Gln851Arg, p.Leu888Pro, p.Arg901Met, p.Leu924Pro, and c.2921+1G>A), occur at sites that are highly conserved among the α subunit paralogs (*ATP1A1 –ATP1A4*) and invariant in other vertebrate *ATP1A3* orthologs (**Figures S1B**). *In silico* pathogenicity predictions for p.Gln851Arg, p.Arg901Met, p.Leu924Pro, and c.2921+1G>A suggest these variants are likely deleterious and disease causing (**Table S1**). Two amino acid substitutions reported here, Leu924Pro and Gln851Arg, were previously associated with EIEE and AHC without MCD(Arystarkhova et al., 2019; Masoud et al., 2017), further supporting the wide spectrum of *ATP1A3* phenotypes resulting from single amino acid variants (**Table S3**). Leu924Pro pathogenicity was also demonstrated to affect ATPase trafficking to the plasma membrane, categorized functionally as loss-of-function (LOF), or viewed as a toxic gain-of-function effect.(Arystarkhova et al., 2019) Together, the conservation of nucleotides and absence of variants in population databases suggests these PMG *ATP1A3* alleles are likely pathogenic.

PMG-associated missense *ATP1A3* variants (p.Gln851Arg, p.Leu888Pro, p.Arg901Met, p.Leu924Pro) cluster within TM7 to TM8 segments. The *de novo ATP1A3* variant in Case A, NM_152296.4:c.2702G>T, p.Arg901Met (R901M), affects an arginine in the extracellular loop connecting transmembrane domains 7 and 8 (TM7-TM8; **Figures 2A, S1A**), in the same segment as a previously described variant (p.Leu888Pro) occurring in an AHC individual with PMG(Gurrieri et al., 2016). Arg901 and Leu888 amino acids are in close proximity to Glu899, Gln904, and Gln905, which interact with the Na,K-ATPase β subunit(Shinoda et al., 2009) **(Figure 2B)**. Case C possesses a *de novo ATP1A3* variant, NM_152296.4:c.2771T>C, (p.Leu924Pro), altering an amino acid in the TM7 helix just upstream of the loop (**Figures 2B**). Case D possesses a *de novo ATP1A3* variant, NM_152296.4:c.2552A>G, p.Gln851Arg, altering an amino acid in TM7 (**Figure 2B**). Although the clustering of these missense variants may be seen as suggesting disruption of specific protein-protein interactions, on the other hand, the *de novo ATP1A3* variant in Case B, NM_152296.4:c.2921+1G>A, abolishes the essential splice donor site of exon 21 **(Figure S1**), and similar *ATP1A3* splice site variants are known to cause AHC(Viollet et al., 2015). Therefore, the PMG-associated variants may all represent LOF, with the more severe phenotype reflecting more complete LOF than for some other ATP1A3-associated conditions.

**Figure 2.**
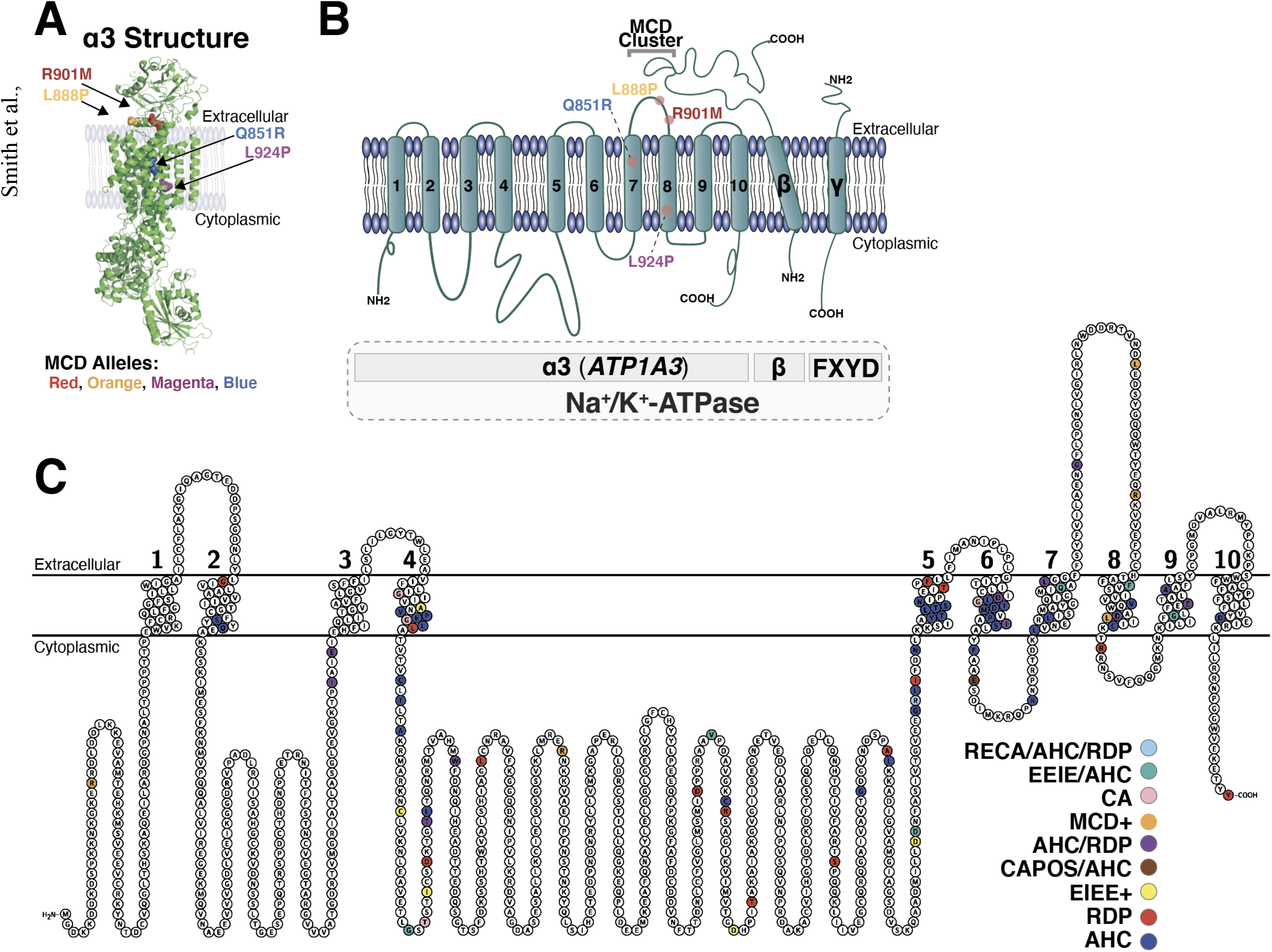
Schematic of *ATP1A3* variants associated with cerebral cortex malformations and other *ATP1A3* related diseases. **(A)** Overview of the α subunit (green) of the Na,K-ATPase with novel PMG alleles mapped (colored amino acids) and previous case report of PMG associated allele (Leu888Pro)(Gurrieri et al., 2016). Red, p.Leu888Pro (L888P); Orange, p.Arg901Met (R901M); Purple, p.Leu924Pro (L924P); Blue, p.Gln851Arg (Q851R). Image generated with PyMOL using Protein Databank 2ZXE(Shinoda et al., 2009). (**B)** Transmembrane topology schematic of Na,K-ATPase for visualization of PMG causing variants, including α, β, and FXYD subunits. We also denote a variant enriched region TM7-TM8, where β subunits interact with the α3 subunit in the extracellular TM7-TM8 segment Glu899, Gln904, and Gln905(Shinoda et al., 2009). (**C)** Overview of genotype-phenotype correlation of *ATP1A3* associated disease single nucleotide variants, phenotypes were grouped by primary symptomatic presentation: including early infantile epileptic encephalopathy (EIEE); alternating hemiplegia of childhood (ACH); cerebellar ataxia, areflexia, pes cavus, optic atrophy, and sensorineural hearing loss (CAPOS); rapid-onset dystonia-parkinsonism (RDP); and childhood onset schizophrenia (COS), and/or a cortical malformation syndrome (MCD+), cerebellar ataxia (CA). Alleles from several sources including(Arystarkhova et al., 2019; Brashear et al., 1993; Holm et al., 2016; Sweadner et al., 2019). See supplemental Figure S7 and Table S3 for complete allele to protein topology breakdown.

### *ATP1A3* mRNA expression in the human fetal neocortex

To investigate the transcriptional trajectory of *ATP1A3* during human brain development underlying the prenatal phenotypes described in this study, we first contrasted the temporal expression patters of *ATP1A1-3* across brain regions, mining bulk neocortical transcriptome data from the Allen Brain Atlas(Jones et al., 2009), ranging from 12 weeks post conception (wpc) to adulthood (40 year-old) (**Figure 3**). We find that prenatal expression of *ATP1A3* is higher than that of its paralogs, and that *ATP1A3* persists postnatally (**Figure 3B**). To gain further insights into the cell-biological mechanisms underlying the developmental cortical phenotypes described in this study, we characterized the patterns of *ATP1A3* expression in the midgestational fetal human neocortex (wpc 19-21). This developmental age offers a uniquely informative window into corticogenesis as many and diverse biological processes – including neurogenesis, neuronal migration and differentiation, and cortical folding – co-occur at this age. At this stage, early-born excitatory neurons (ENs) have settled and started to differentiate within the innermost strata of the cortical plate (CP) (prospective layers 6a-5), while later-born ENs progressively migrate past their predecessors to occupy the most superficial aspect of the CP (prospective layers 4-2)(Miller et al., 2014). A large fraction of upper-layer ENs are still being generated in the germinal zones, while others – en route to the CP – are witheld within the subplate (SP), a transient developmental layer crucial for corticogenesis, where ENs are primed by thalamic input and form transient synaptic connections with one another and with resident SP neurons (prospective layer 6b)(Luhmann et al., 2018).

**Figure 3.**
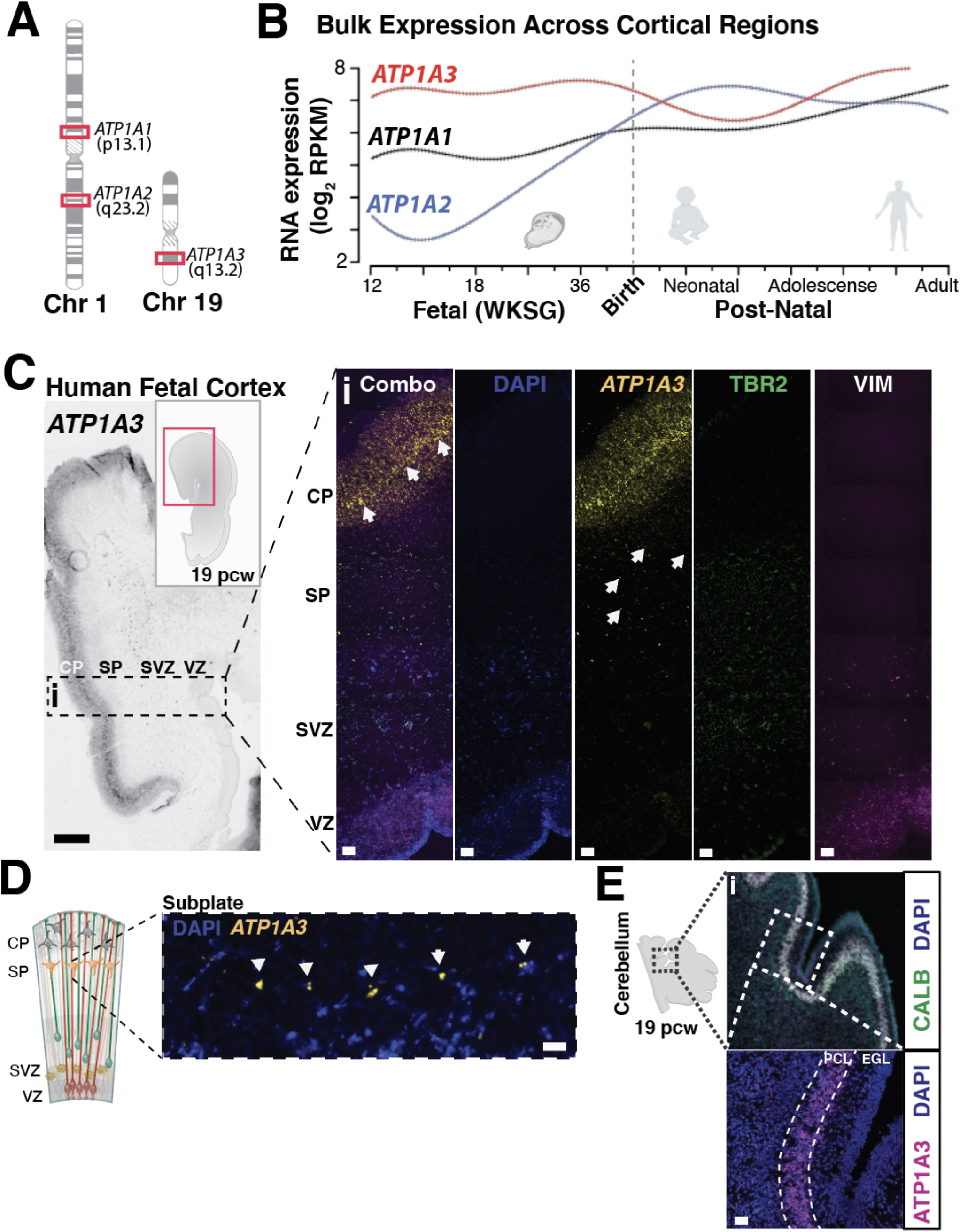
*ATP1A3* is differentially enriched to the human fetal period and localized to subplate and deep-layer neurons. (**A)** Chromosome locations for CNS enriched Na^+^,K^+^-ATPase alpha isoforms, *ATP1A1* and *ATP1A2* (Chr1), and *ATP1A3* (Chr19). (**B)** RNA transcriptome analysis of bulk brain regions revealed *ATP1A3* transcripts are high during fetal gestational and persist postnatally. CNS expressed paralogous *ATP1A1* and *ATP1A2* paralogs show relative expression pattern changes during development, while *ATP1A4* is not expressed significantly in the brain. Raw transcriptome data for (B) from Allen Brain Atlas, presented as log2 RPKM (reads per kilobase per million) values and a polynomial fit to average across time points(Jones et al., 2009). (**C)** *Left, ATP1A3* mRNA chromogenic *in situ* hybridization performed on 19 weeks post conception (pcw) coronal brain sections demonstrate highest *ATP1A3* expression in the human cortical plate (CP), and subplate region (SP). Scale bar left, 500μm. *Right*, Corresponding fluorescence imaging of 19 wpc fetal brain with cell type specific markers for intermediate progenitors (TBR2) and neural progenitors (vimentin, VIM) demonstrates *ATP1A3* transcripts are not present within cells of the sub-ventricular zone (SVZ) and ventricular zone (VZ). Arrows indicate high expressing neurons in deep cortical layers and subplate region. (**D**) Magnified image from (C) depicting high *ATP1A3* expressing subplate layer cells. (**E**) *Left*, Schematic of developing sagittal fetal cerebellum tissue section at 19 wpc demonstrates enrichment of *ATP1A3* in Purkinje cell layer. (*i*) *ATP1A3* mRNA fluorescence in situ at 19 wpc demonstrates highest expression in the Purkinje cell layer, colocalizing with Purkinje cell marker calbindin (CALB1, green). (ii), Zoomed fluorescence image, including labeling of external granule layer. Scale bar left, 50μm. DAPI stain for nuclei in blue. Scale bar 25μm. VZ, ventricular zone; SVZ, subventricular zone; IMZ, intermediate zone; SP, subplate; CP, cortical plate; PCL, purkinje cell layer; EGL, external granule layer.

To investigate *ATP1A3* expression in its tissue context, we perfomed RNA *in situ hybridization* (ISH) in a coronal cross-section of fetal neocortex at 19 wpc. *ATP1A3* mRNA was virtually undetectable in the germinal zones (ventricular and subventricular zone, VZ and SVZ) but abundant in postmitotic neuronal layers, with highest expression in the inner CP and SP (**Figure 3C & D**). Multiplexed fluorescent ISH analysis of *ATP1A3* with markers of two populations of cortical neural progentiors cells, radial glia and intermediate progentior cells, confirmed *ATP1A3* is not expressed in these dividing cells or their immidiate progeny in either the VZ or the SVZ (**Figure 3C**). In addition, since *ATP1A3*-associated diseases display a range of cerebellar deficits(Brashear et al., 1993; Sweadner et al., 2016; Uchitel et al., 2019), we also evaluated *ATP1A3* expression in the fetal cerebellum (19 wpc), demonstrating an exclusive enrichment of *ATP1A3* within the Purkinje cell layer (**Figure 3E**).

### Fetal neocortex single cell analysis demonstrates robust *ATP1A3* enrichment to resident subplate neurons and rostral neocortical *ATP1A3* enrichment

Despite the high degree of compartmentalization of cell types into laminae, biological processes in the developmental neocortex are highly dynamic, and cellular compartments are heterogeneous in cell-type composition. Single-cell RNA sequencing enables large-scale, simultaneous comparisons of gene expression across thousands of individual cells, thus bridging functional data from human genetics, immunohistochemistry, and cell-type-specific biology (Macosko et al., 2015). To this end, we performed Drop-seq on 125,942 individual cells from a 21 wpc human neococrtex **(Figure 4A & S2A**). Given the general trend of perisylvian to frontal cortex localization of the PMG phenotypes described in this study, we sought to investigate *ATP1A3* expression across presumptive neocortical areal domains. To build our single-cell expression dataset, we collected cells from 11 adjecent sections spanning the dorso-medial and rostro-caudal axes of the neocortex, validating and refining the positional identity of these serial samples using the graded expression of cortical patterning genes as benchmarks (**Figure 4B & S2**)(Sun et al., 2005). Using unsupervised statistical methods (see materials and methods), cells were partitioned into four transcriptionally distinct clusters, corresponding to the major neural cell classes in the midgestational neocortex: excitatory neurons (EN), inhibitory neurons (IN), neural progenitor cells (NPCs), and glia (**Figure 4C & S2C**). We excluded from this analyisis all non-neural cell types as our scope of inquiry was specifically regarding *ATP1A3* dysfunction in neurons, and non-neural expression of *ATP1A3* is limited (**Figure 3B & S2**). Within each of the main neuronal cell classes we could further resolve transcriptionally distinct cell subtypes and states, which we interpreted by the distinctive enrichement of marker genes in each group (**Figure S2C**). Among ENs, consistent with the developmental landscape of mid-gestational corticogenesis described above, we could identify newborn and migratory upper-layer ENs at different stages of differentiation (clusters EN.0-3), and post-migratory deep-layer ENs expressing marker genes of maturing layer 6-5, including markers of resident SP neurons (EN-4, **Figure 4C**). Across cortical regions, *ATP1A3* expression was virtually undetectable in the NPC clusters, low in INs and glia, more highly expressed in ENs (**Figure 4C, S2C & D**), and – among ENs – highest in the EN-4 cluster.

**Fig 4.**
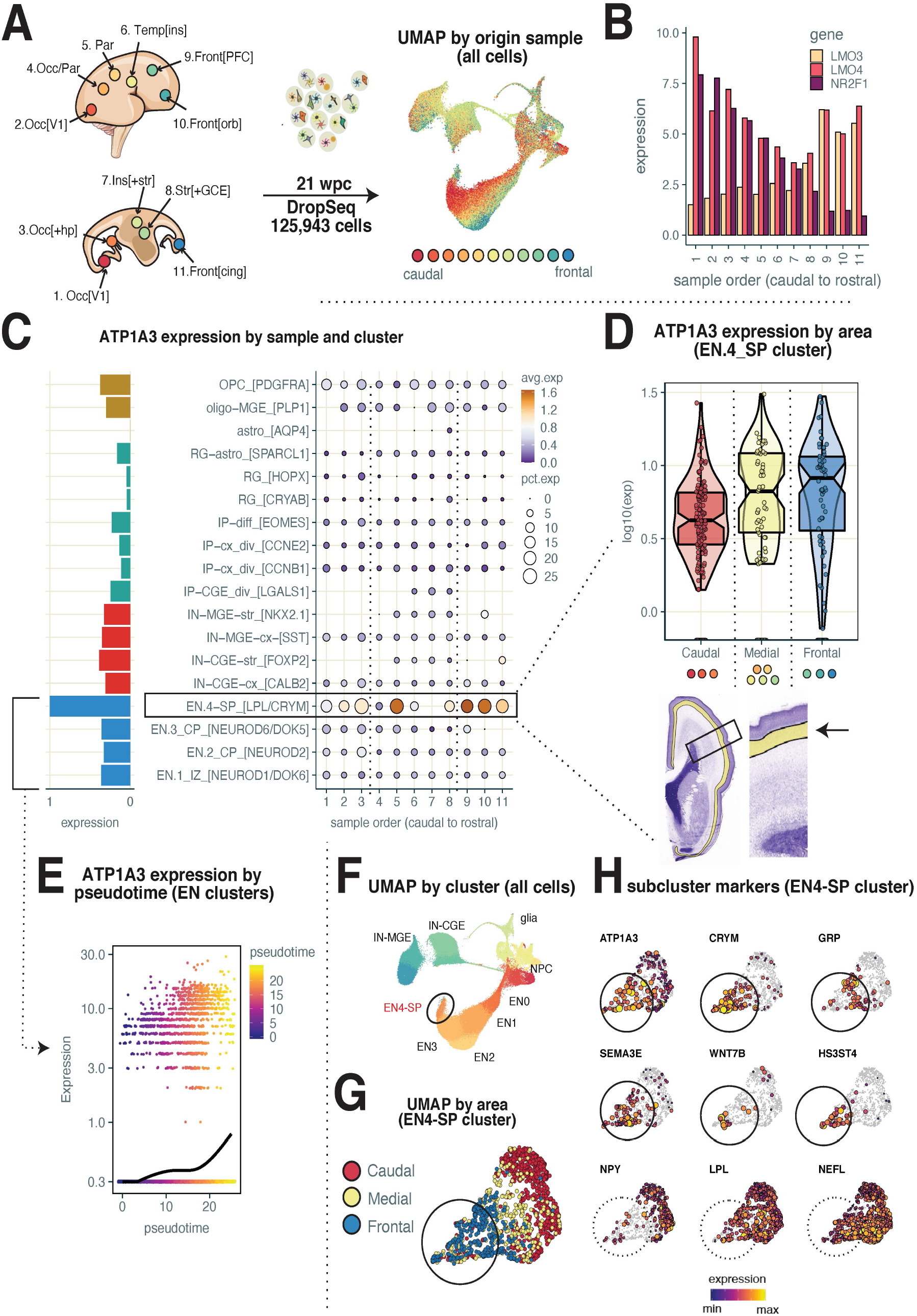
Single cell *ATP1A3* expression atlas in the developing human cerebral cortex. (**A**) *Left:* Schematic showing a 21 post conception weeks (pcw) forebrain and location of regions sampled (for anatomical detail see fig. S2A). *Top:* dorsolateral view. *Bottom:* sagittal view. *Right:* Uniform Manifold Approximation and Projection (UMAP) of 125,943 single cells profiled by DropSeq. Each dot represents a cell, color-coded by sample of origin. Key: blue-to-red: frontal-to-caudal. Occ: Occipital; Par: Parietal; Temp: Temporal; Ins: Insula; Orb: Orbital; Str: Striatum; PFC: Prefrontal Cortex; V1: Primary Visual Cortex; Hp: Hippocampus; CGE: Caudal Ganglionic Eminence; MGE: Medial Ganglionic Eminence (**B**) Graded expression of areal marker genes *LMO3, LMO4* and *NR2F1* (COUP-TF1) across samples ordered caudal-to-rostral (x-axis). Gene expression (y-axis) was normalized by cell (divided by total UMIs/cell x 100k), then the mean value for each gene was aggregated by sample. Notice the opposite gradient of *LMO3* and *NR2F1* expression, and the U-shaped pattern of LMO4 expression along the rostro-caudal axis. See fig. S2B for additional areal markers. (**C**) *Left:* Histogram showing relative expression of *ATP1A3* (x-axis) across cell clusters (y-axis). Notice enriched expression of *ATP1A3* in cluster EN.4-SP, containing subplate (SP) EN neurons (See **Fig. SC** and **H** for SP markers enriched in this cluster). Mean *ATP1A3* expression was aggregated by cluster, then rescaled from 0 to 1. Cell clusters are color-coded by cell type (blue: excitatory neurons (EN); gold: glia; blue: inhibitory neurons (IN); red: neural progenitor cells (NPC). See fig.SB for cluster markers and assignments. *Right:* Dot-plot showing *ATP1A3* expression across clusters split by areas of origin (x-axis; sample 1-11, ordered from caudal to rostral). Color scale codes for mean expression by group; size of the dots codes for percentage of cells expressing *ATP1A3* in each group. (**D**) *Top*, Violin-Boxplot showing differential expression of *ATP1A3* (y-axis) in the EN.4-SP cluster across three main cortical partitions (x-axis; Caudal: samples 1-3 (occipital cortex); Medial: samples 4-8 (including parietal and temporal cortex, and subcortical structures); Frontal: samples 9-11 (frontal cortex). Dots represent individual cells. Violins show probability density distributions; boxes show interquantile ranges; notches show confidence intervals around the median (horizontal lines). Outliers were trimmed. Mean expression of *ATP1A3* was aggregated by region, then log10-transformed. Notice the highest expression of *ATP1A3* in cells sampled from frontal cortex, and lowest expression in cells sampled from caudal cortex. *Bottom:* Nissl-stained frontal coronal section of a 21 pcw forebrain highlighting anatomical position of SP. Source: Atlas of the Developing Human Brain, *BrainSpan* (www.brainspan.org), Reference Atlas. (**E**) *ATP1A3* expression across cells in the EN clusters (color-coded in blue in C, left) ordered by pseudotime. Dots represent individual cells. Y-axis: log-transformed, scaled expression of *ATP1A3* by cell. X-axis: pseudotime score (color-coded) for each cell, calculated using the Monocle3 algorithm. Trend-line shows the increase of *ATP1A3* expression as function of pseudotime, calculated by fitting a quasipoisson model to the data. (**F**) UMAP of all profiled cells color-coded by cluster (see C, left). The EN.4-SP cluster is highlighted. (**G**) UMAP of cells in the EN.4-SP cluster (highlighted in F) color-coded by rostral, medial and caudal anatomical partition of origin (see also D, top). Notice that cells segregate by origin. (**H**) Relative expression of areal marker genes in the EN.4-SP cluster (see G). Relative gene expression is coded by color and size. Notice that *ATP1A3* is enriched in cells expressing SP markers (*CRYM, WNT7B, HS3ST4*) and frontal cortex markers (*GRP, SEMA3E*), but not caudal cortex markers (NPY, LPL, NEFL).

The EN-4 cluster was composed of cells sourcing from all cortical regions, and was marked by enriched expression of SP-specific genes (*CRYM* and *LPL*, **Figure 4C**), as well as by the expression of genes associated with neuronal maturation (e.g. axonal genes *NEFL/NEFM* and synaptic genes *SEMA3E*). This is consistent with the notion that the earliest-born neurons (a subset of which will persist in L6b in the adult cortex) settle and mature early in the developing SP(Hoerder-Suabedissen and Molnár, 2013). Within this cluster, *ATP1A3* appeared to be differentially expressed across cortical region, with highest expression levels found in cells from frontal samples, intermediate levels in cells from medial samples, and lowest levels in cells sampled from more caudal cortical regions (**Figure 4C & S2A**). In agreement with this, expression of *ATP1A3* in the EN-4 cluster was found to be enriched in neurons co-expressing SP markers and frontal cortex markers, but much less so in neurons expressing caudal cortex markers (**Figure 4D, S2 & S3)**.

Cortical development proceeds along a rostral-to-caudal maturation gradient, with asyncronous growth patterns accounting for the differential expansion of frontal and caudal cortical regions(Rakic, 1974). This implies that, at any given time, neurons in frontal regions are on average more mature than their caudal counterparts. To investigate whether the observed *ATP1A3* enrichment in the EN-4 cluster might reflect a differential maturation state of EN neurons along the rostrocaudal axis, we performed pseudotime analysis of *ATP1A3* across all ENs. To do this, we used the Monocle3 algorithm, an established analytical tool allowing single-cell transcripomes to be ordered and connected along a branched topology based on the differential expression of variable genes(Cao et al., 2019). By selecting as an “origin” cell, this algorithm allows projection of a “pseudo-differentiation” trajectory onto the network topology(Camp et al., 2015). This analysis revelaed a positive correlation bewteen *ATP1A3* expression and EN differentiation, suggesting *ATP1A3* may be required during neuronal differentiation, as neurons transition to a mature state, most readily observed in resident subplate neurons (**Figure 4E & F).**

Finally, to identify potential isoform-specific and cell-type specific vulnerabilities to *ATP1A1-3* variants during corticogenesis, we compared the expression patterns of the *ATP1A1-3* paralogs across cell clusters (**Figure S3A**). We find that expression of *ATP1A2* is restricted to NPCs (and specifically to radial glia), and is virtually undetectable in neurons – in an almost mutually-exclusive pattern to *ATP1A3*. *ATP1A1* expression more closely mimicked *ATP1A3* expression, albeit its expression in NPCs was higher, and partially overlapped *ATP1A2* (**Figure S3A**). Accordingly, pairwise correlation analyis of the three isoforms across clusters revealed that expression of *ATP1A1* and *ATP1A3* correlates more strongly with the EN-4-SP cluster, while expression of *ATP1A1* and *ATP1A2* correlates more highly with NPC clusters (**Figure S3B**). Across all cells, pairwise correlation analysis indicated a positive correlation, though moderate, between *ATP1A1* and *ATP1A3*, but weak-to-no correlation between *ATP1A1* and *ATP1A2*, and between *ATP1A2* and *ATP1A3* (**Figure S3C**). Together, this analysis suggests that neurons transitioning within the fetal SP may be particularaly susceptible to *ATP1A3* variants, with potential broad consequences for cortical lamination. Moreover, the single cell variable expression of ATPase isoforms could underlie compensation or vulnerbility mechanisms for disease haploinsufficiency or differental penetrance within specific cell populations.

### *ATP1A3* enrichment in parvalbumin interneurons and excitatory neurons in infant neocortex

Given that *ATP1A3* expression persists in the postnatal neocortex (**Figure 3B**), and that several known *ATP1A3* mutations show postnatal onset of symptoms(Brashear et al., 1993; Smedemark-Margulies et al., 2016), we profiled *ATP1A3* expression in the infant neocortex. Using Drop-seq, we profiled 51,878 nuclei from a ~8-month-old human cortex, sampled from prefrontal, temporal, occipital, and parietal lobes (**Figure 5A**). Unsupervised hierarchical clustering sorted nuclear transcriptomes into 3 major classes – ENs, INs, and glia – subdivided into 24 transcriptionally distinct cell classes (**Figure 5A** & **S4**). Within the EN clade, clusters stratified largely by layer and neuronal subtype, rather than cortical area, with several clusters enriched in gene markers of deep layer 5 and 6 (**Figure 5B,** See supplemental **S4** for full clading). Within the IN clade, cells stratified by developmental origin (Caudal or Medial Ganglionic Eminence) and major “cardinal types”, according to previous studies(Mayer et al., 2018). In the infant neocortex, like in the fetal neocorex, *ATP1A3* was also enriched in neurons compared to glia, but unlike in the fetal dataset, *ATP1A3* was more highly expressed in INs than in ENs **(Figure 5B & S5)**. Among neuronal clusters, and across cortical regions, *ATP1A3* expression was fairly variable, with lowest expression in V1/occipital cortex cells, and wide range of expression levels within deep and upper layer EN clusters (**Figure 5B**). No specific enrichment was found in L6b ENs (i.e. the postnatal remnant of the fetal SP layer), suggesting that fetal enrichment of *ATP1A3* in the SP is transient, with maturation and migration of cells continuing to populate the upper layers. Of note, the lack of specific enrichment in postnatal SP may reflect the fact that, unlike the fetal dataset, the postnatal dataset was built from nuclear RNA rather than whole cell RNA, so that some transcripts may be under-detected, primarily those localized primarily in the somato-dendritic compartment.

**Figure 5.**
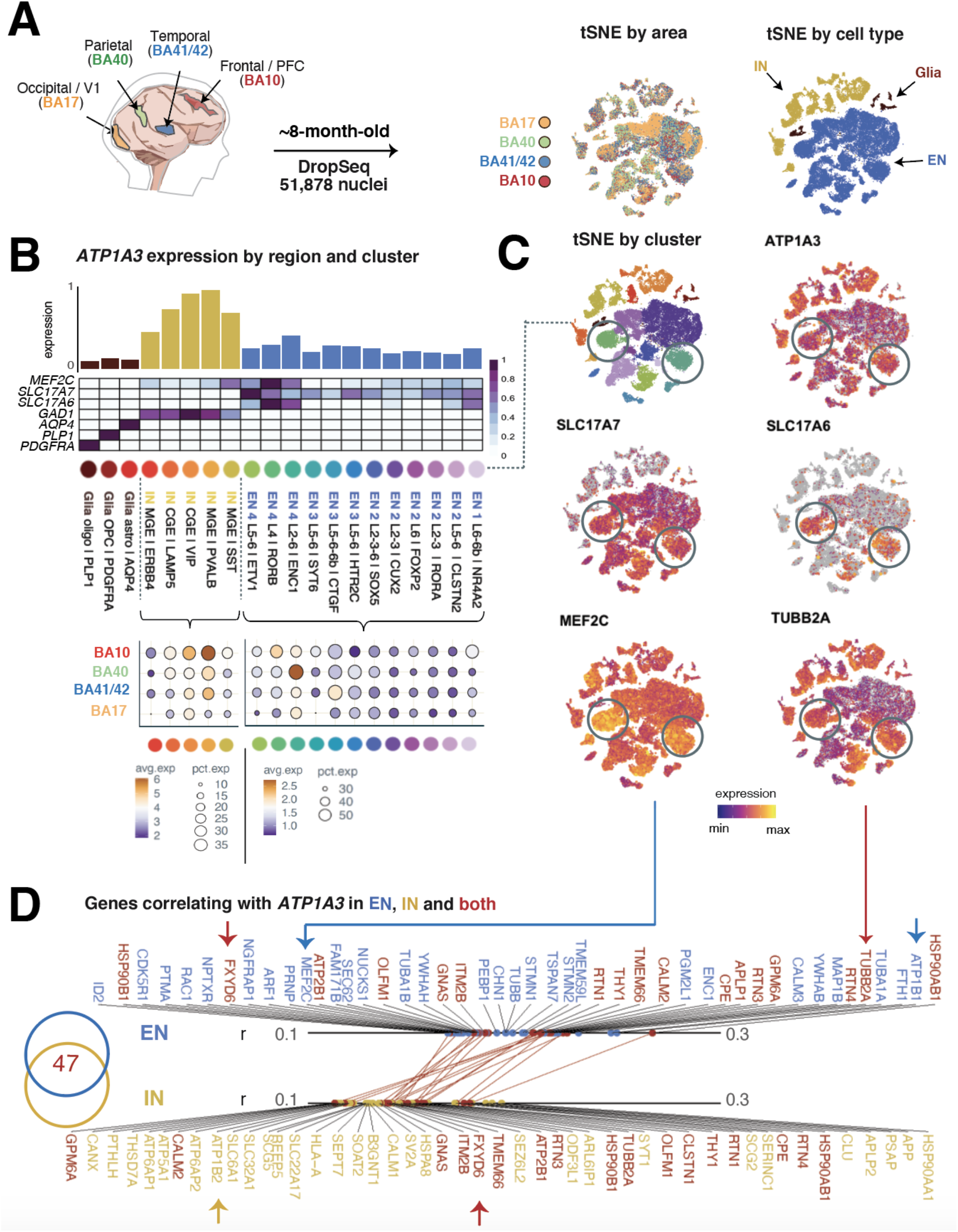
Single-cell analysis of *ATP1A3* expression in the infant human neocortex. (**A**) *Left*, Schematic showing an 8-month-old neocortex and location of regions sampled. BA: Brodmann area; PFC: Prefrontal Cortex; V1: Primary Visual Cortex. *Right*, T-distributed Stochastic Neighbor Embedding (tSNE) of 51,878 single nuclei profiled by DropSeq. Each dot represents a nucleus, color-coded by cluster (*left*), cell type (*middle*) and origin sample (*right*). EN: excitatory neurons; IN: inhibitory neurons. See C for cluster and cell-class assignments. (**B**) *Top*, dendrogram summarizing hierarchical clustering of data. Clusters are color-coded as in A, left). See fig.S4 for cluster markers and assignments. *Middle*, bar graph showing relative expression of *ATP1A3* (y-axis) across cell clusters (x-axis). Mean expression was aggregated by cluster, then rescaled from 0 to 1 across all clusters. Data are color-coded by cell type (as in A, middle – blue: EN; gold: IN; brown: glia). *Bottom*, Dot-plot showing *ATP1A3* expression across the IN and EN clusters (x-axis) split by areas of origin (y-axis). Color scale codes for mean expression by group; size of the dots codes for percentage of cells expressing *ATP1A3* in each group. *Right*, tSNE plots showing specific enrichment of cortical marker genes including genes associated with early activity in the developing cortex (*MEFC2, SLC17A7* and *SLC17A6* (VGLU1 and VGLU2), and MCD associated gene TUBB2A. (**D**) Venn diagram showing intersection between the top-100 genes correlating with *ATP1A3* in the EN cluster (blue) and IN cluster (gold). 47/100 genes are shared between the two groups. Top 50 genes correlating with *ATP1A3* in the EN and IN clusters are shown. Dots indicate Pearson’s R for each gene in the EN cluster (*top*; blue) and IN cluster (*bottom*; gold). Genes found in both groups are indicated in red, and the respective correlation coefficients are connected by red lines.

Across INs clusters, expression of *ATP1A3* was highest in cells sampled from the frontal lobe (prefrontal BA10), and more strikingly so within Parvalbumin (PV^+^) INs (**Figure 5B & S4C**). This differential expression pattern raised the hypothesis that *ATP1A3* may be required dynamically across neuronal types, cell-biological processes, and across developmental ages. To gain insights into the nature of the cell-biological signature associated with *ATP1A3* expression, we compared the top-100 genes correlating with *ATP1A3* in EN and IN clusters. Interestingly, we found that 50/100 *ATP1A3-correlating* genes were shared across cell types, suggesting that *ATP1A3* partakes in similar transcriptional programs, and thus potentially similar function, in different neuronal types. In both EN and IN, among the top genes correlating with *ATP1A3*, was *FXDY6*, the FXYD subunit of the ATPase complex **(Figure 5D)**. We did find, however, that the ATPase β subunit beta subunit *ATP1B1* correlated with *ATP1A3* only in ENs, and *ATP1B2* in INs, suggesting that, at least in part, *ATP1A3* may form cell-type specific ATPase complexes in different neuronal classes **(Figure 5D)**. Gene Ontology Enrichment analysis of the top 100-genes most strongly correlated with *ATP1A3* in both ENs and INs revealed significant over-representation of gene products associated with transmembrane ionic transport and the somatodendritic compartment, including synaptic vesicles **(Table S2)**. This is consistent with the established role of *ATP1A3* in synaptic transmission and membrane potential, and suggests *ATP1A3* may partake in a shared transcriptional network for which *ATP1A3* variants may perturb aspects of neuronal physiology central to many cell types.

Given the association of *ATP1A3* pathological variants with phenotypes affecting higher-order cognitive functions with no clear counterpart in animal models, it was of interest to investigate the degree to which *ATP1A3* enrichment in INs may be conserved among humans, non-human primates, and rodents. We took advantage of a recently published single-cell RNAseq dataset profiling gene expression in homologous IN types in humans, non-human primates (macaques and marmosets) and mice(Krienen et al., 2019). We found that relative *ATP1A3* expression levels across IN types, both in the neocortex and in the striatum, are largely conserved across species, with highest expression in the fast spiking PV^+^ INs (Figure S6). This suggests that the observed differential expression of *ATP1A3* across neuronal types is genetically encoded and evolutionarily conserved.

## DISCUSSION

Mutations in *ATP1A3*, the gene encoding the Na,K-ATPase *a3* catalytic subunit, have been associated with a wide spectrum of neurological diseases affecting childhood development, ranging in onset age from early infancy to late adolescence into adulthood. Here, we describe *ATP1A3* pathogenic variants in individuals presenting with PMG, a severe prenatal malformation of the cerebral cortex characterized by abnormal gyrification and laminar organization. These case studies extend the known phenotypic range of *ATP1A3-related* diseases, and provide new insights into the molecular pathology of PMG and PMG-associated developmental channelopathies. Furthermore, by compiling a single-cell atlas of *ATP1A3* expression in the fetal and early postnatal neocortex, we revealed cell-type-specific *ATP1A3* expression in the fetal and infant human brain, establishing a spatiotemporal roadmap for future genotype-phenotype associations studies, and studies of *ATP1A3* function in the subplate and cortical interneurons.

*ATP1A3* associated conditions comprise a broad spectrum of overlapping phenotypes, including the blending of encephalopathies with more paroxysmal phenotypes, with a complex genotype-phenotype correlation. Further confounding the genotype to phenotype relationship, identical amino acid substitutions can impart variable presentations (**Table S3**); for example, Leu924Pro and Gln851Arg reported here have been previously associated with EIEE and AHC without MCDs(Arystarkhova et al., 2019; Masoud et al., 2017), but with postnatal microcephaly (Leu924Pro), suggesting pathogenic susceptibility differs between individuals. Differential penetrance between individuals may reflect α3 ATPase expression levels, competition between disease and wild-type α3 subunits for utilization within Na/K-ATPases(Arystarkhova et al., 2019), and as described in this study, variable expression of α3 (and coexpression with other α and β subunits) within different cell populations (**Figure 4D, S3 & S5**). Individuals with heterozygous *ATP1A3* variants presented here with PMG also exhibited a range of phenotype severity, from unilateral to extensive bilateral PMG, variable degrees of developmental delay, postnatal microcephaly, ataxia-like phenotypes and EIEE. *ATP1A3’s* wide phenotypic spectrum is similar to known glutamate and sodium channel “developmental channelopathies”, in which a subset of cases present with EIEE and developmental delay, both with and without MCDs and postnatal microcephaly(Smith and Walsh, 2020). Taken together, *ATP1A3* PMG-associated variants likely represent a continuum of LOF features, with the more severe phenotype reflecting more complete LOF than for some other *ATP1A3*-associated conditions. In a future opportunity to examine cortical tissue of PMG associated with *ATP1A3* variants, it would be useful to document cortical architecture, including whether gaps occur in the pia mater and glia limitans. This anatomical defect permits continuity between the molecular zones of adjacent microgyri for synaptic excitatory short-circuitry leading to epilepsy, as with other genetic mutations in PMG(Jansen et al., 2016).

Functional analysis of *ATP1A3* variants associated with postnatal diseases are largely described as LOF; including decreased membrane localization in Leu924Pro, causing toxic effects within cells(Arystarkhova et al., 2019). LOF mechanisms described for *ATP1A3* disease-causing variants include changes in Na^+^ affinity, impaired conformational changes, voltage-dependence shifts, and disruption of protein expression(Arystarkhova et al., 2019; Heinzen et al., 2014; M. Li et al., 2015). Since the Na,K-ATPase exports Na^+^, and elevated Na^+^ can be deleterious to cellular processes(Azarias et al., 2013; Toustrup-Jensen et al., 2014), α3 dysfunction that results in a shift in Na^+^ affinity could trigger a known developmental pathophysiology associated with increased intracellular Na^+^ concentration, leading to PMG (Smith et al., 2018; Smith and Walsh, 2020; Zaman et al., 2020). While some *ATP1A1* variants have been described to increase inward leak of Na^+^(Azizan et al., 2013; Kaneko et al., 2014), similar pathogenic gain of function effects have not been described in *ATP1A3* related diseases. In addition to acute ATPase pump dysfunction, PMG pathophysiology could also reflect an accelerated activation of toxic downstream mechanisms that underlie postnatal *ATP1A3*-related disorders causing neurodegeneration or cell death, including postnatal microcephaly(Paciorkowski et al., 2015), regional neuronal loss(Oblak et al., 2014), and cerebellar hypoplasia(Sweadner et al., 2016; Uchitel et al., 2019). Moreover, two of the PMG-causing *ATP1A3* variants we describe are located in the TM7-TM8 extracellular segment of the α3 that interacts with the β-subunit(Shinoda et al., 2009) and Leu924Pro demonstrates β subunit associated LOF membrane localization properties(Arystarkhova et al., 2019), suggesting the final formation of α-β complex could potentially underlie the pathophysiology. The β-subunit supports localization of the α-subunit to the plasma membrane and modulates affinity of Na^+^ and K^+^ for the ATPase(Blanco et al., 1998). Moreover, α- and β-subunits can trigger several intracellular signaling pathways associated with MCDs, including MAPK and AKT signaling via phosphoinositide 3-kinase (PI3K) and EGFR pathways, resulting in changes in cell polarity, growth, motility and gene expression(Z. Li and Langhans, 2015).

During embryonic brain development, bioelectric cellular properties are required for the propagation of key organizing signals, directly implicating ion channels and pumps in a variety of morphogenetic processes(Levin et al., 2017). For instance, in mice and ferrets, experimental modulations in membrane conductance and Na+-dependent excitation have been linked to neuronal fate specification and directed migration(Hurni et al., 2017; LoTurco et al., 1995; Smith et al., 2018; Vitali et al., 2018). The fetal SP represents a hub for the spatiotemporal integration and transduction of signals from many cellular sources. Early electrical activity patterns in the SP accompany the dynamic formation of transient synaptic networks between resident SP neurons and migratory ENs and INs en route to the CP, ascending radial glia processes, and ingressing thalamic fibers^23,46^. Our finding that (i) *ATP1A3*-expressing cells are located in the developing SP (ISH, Fig 3D), and (ii) co-express SP-specific marker genes (DropSeq, Fig 4 and S2), is therefore consistent with a role for *ATP1A3* in regulating electric activity within this compartment. This is consistent with the notion that human SP neurons possess large Na^+^ currents with prolonged depolarized states(Moore et al., 2009), and therefore require a stout mechanism to recover Na^+^ and K^+^ electrochemical gradients following excitation, a known key function of the α3 ATPase in postnatal neurons(Kim et al., 2004; Picton et al., 2017; Vaillend et al., 2002). In addition, the finding that *ATP1A3-expressing* cells are also found in the inner aspect of the CP (ISH, Fig 2C), express L6-5 EN markers (DropSeq, Fig4C), together with the rostral-to-caudal gradient of *ATP1A3* expression and positive correlation with neuronal pseudo-age, (Monocle, Fig 4F), all are consistent with a role of *ATP1A3* in the maturation of early-born deep-layer ENs.

The observed differential prenatal expression of Na^+^,K^+^,ATPase isoforms parallels the enrichment pattern reported for specific glutamate and sodium channel subtypes implicated in developmental channelopathies. These glutamate and sodium channel diseases maintain robust RNA expression of developmental subtypes; specific channel subtypes are utilized for developmental electrical functions (*SCN3A, GRIN2B*), whereas others are used for adult electrical functions (*SCN1A, GRIN2A*)(Smith and Walsh, 2020). The α3 isoform may have a similar specialized function in fetal brain neurons compared to α1 isoforms. For example, Na^+^ affinity is lower in α3 than α1 isoforms(Azarias et al., 2013; Dobretsov and Stimers, 2005), enabling α3 containing ATPase pumps to respond to larger Na^+^ influxes. Lower Na^+^ affinity provides a wider range to buffer aberrant Na^+^ flux, therefore could provide an evolutionarily advantageous neuroprotective effect in cells during development. Known developmental channelopathy genes also maintain a robust cell type and subtype expression profiles as *ATP1A3;* for example, *SCN3A* and *GRIN2B* are enriched in excitatory neurons and some progenitor types(Bagasrawala et al., 2017; Smith et al., 2018; Smith and Walsh, 2020), before transitioning to mostly *GRIN2A*-containing glutamate receptors and *SCN1A* in the postnatal cortex(Bagasrawala et al., 2017; Smith and Walsh, 2020). Intriguingly, the observed sc-RNA-seq *ATP1A2* enrichment in NPCs could offer a pathological basis for *ATP1A2* variants associated with fetal or early life demise, including microcephaly(Chatron et al., 2019; Monteiro et al., 2019).

In the postnatal cortex, the observed *ATP1A3* enrichment in interneurons likely supports physiological processes necessary for maintaining inhibitory tone of cortical circuits(Z. Li and Langhans, 2015), a proposed mechanism underling EIEE, or perhaps AHC. We were surprised to find that even after 90 million years of divergence, the human and rodent postnatal cortex possess similar single-cell distribution of *ATP1A3*, most notably in PV^+^ INs, underscoring the utility of mice as a model for studying excitatory/inhibitory balance(Bøttger et al., 2011), especially in the context of PV IN loss relating to postnatal brain disorders(Hunanyan et al., 2018). On the other hand, while *ATP1A3* expression in INs appears to be conserved, differences in circuits and connectivity, and how interneurons shape gene expression differentially could offer pathological differences between species. For example, the observed layer specific rostral enrichment of *ATP1A3* could contribute an early susceptibility for *ATP1A3* variants associated childhood onset schizophrenia(Chaumette et al., 2018; Smedemark-Margulies et al., 2016). Intriguingly, the identified *ATP1A3* enriched cell types in the pre- and postnatal cortex (subplate and PV^+^ neurons), and Purkinje cells of the cerebellum, are all characterized by their robust firing properties(Moore et al., 2009; Tremblay et al., 2016). Within these cell types with dynamic physiological needs, the low affinity α3 would help maintain ionic gradients following excitation and suggests neurons with higher neurophysiological demands would be most susceptible to Na^+^,K^+^,ATPase dysfunction. Additionally, even within the cortical pyramidal neuron population, differential Na^+^,K^+^,ATPase activity levels exists (Anderson et al., 2010). Therefore, *ATP1A3* PMG-associated variants are likely LOF (affecting levels of Na^+^ accumulation in cells), with severe phenotypes reflecting more complete LOF effects and toxicity within more vulnerable cell types.

## Supporting information

Supplemental Results

## ACKNOWLEDGMENTS

We are grateful to the participant families presented here and thank members of the Walsh lab for helpful discussions and Johnathan Hecht for assistance with human tissue. Human tissue was obtained from the NIH Neurobiobank at the University of Maryland, Baltimore, MD. This work was supported by NIH F32NS100033801 and K99NS112604 (R.S.S.), NIH T32GM007753 (S.K.A.), Paul G. Allen Frontiers Program and NIH R01NS032457 and R01NS035129 (C.A.W.), and BCH IDDRC U54 HD090255. This work was also supported by the Broad Center for Mendelian Genomics (UM1 HG008900) funded by the National Human Genome Research Institute (NHGRI) and the Tommy Fuss Foundation (R.S.S and J.G-H.). C.A.W. is an Investigator of the Howard Hughes Medical Institute.

## AUTHOR CONTRIBUTIONS

C.A.W., A.J.B., R.S.S., S.K.A., F.P.V., E.M., L.F.S. and L.B. collected and evaluated clinical data; J.E.N. coordinated clinical data; R.S.S. performed and analyzed tissue experiments; M.F. and R.S.S. performed cortical dissections; M.G., C.D.M., N.R. and M.F. generated Drop-Seq libraries; M.F. analyzed Drop-Seq data; R.S.S., M.F., J.E.N., S.K.A. and C.A.W. wrote the manuscript, with contributions from all co-authors. We thank Kathleen Sweadner for productive discussion. The authors declare no competing financial interests.

## GLOSSARY

ATP1A3: Na^+^,K^+^-ATPase alpha 3 subunit
PMG: Polymicrogyria (an overfolded cerebral cortex)
MCD: Malformation of cortical development
EN: Excitatory neuron
IN: Inhibitory neuron
NPC: Neural progenitor cell
SP: Subplate
Vm: Resting membrane potential
WPC: Weeks post conception
AHC: Alternating hemiplegia of childhood
RDP: Rapid-onset dystonia-parkinsonism
CNS: Central Nervous System

