## Supplemental Results for "Role for the Na^+^/K^+^ ATPase pump alpha 3 (*ATP1A3*) subunit in folding and lamination of the human neocortex"

### Folding and lamination of the human neocortex depend on the sodium potassium pump alpha 3 (*ATP1A3*) subunit

### EXPERIMENTAL DETAILS

#### Human subjects and samples

Individuals presented herein were identified and evaluated in a clinical setting, and biological samples collected after obtaining written informed clinical and/or research consent. Human subject research was conducted according to protocols approved by the institutional review boards of Boston Children's Hospital, Beth Israel Deaconess Medical Center and the Mayo Clinic. Fetal brain tissue was received after release from clinical pathology, with a maximum post-mortem interval of 4 h. Cases with known anomalies were excluded. Tissue was transported in Hibernate-E medium (Thermo Fisher) on ice to either the Walsh laboratory or McCarroll laboratory for downstream processing. The neonatal brain sample was obtained from the University of Maryland Brain and Tissue Bank of the NIH NeuroBioBank (sample number UMBN 5817) and stored at -80C until further processing.

#### Human genetics sequencing and analysis

For Case A and B, whole exome sequencing and data processing were performed by the Genomics Platform at the Broad Institute of MIT and Harvard. Libraries from DNA samples (>250ng of DNA, at >2 ng/ul) the Human Core Exome Kit from Twist Biosciences was used to capture target regions (~38 Mb target), and sequencing was performed on the Illumina NovaSeq 6000 (150 bp paired reads) to cover >96% of targets at 20x and a mean target coverage of >100x. Sample identity quality assurance checks were performed on each sample. The exome sequencing data was de-multiplexed and each sample's sequence data were aggregated into a single Picard BAM file. Exome sequencing data was processed through a pipeline based on Picard, using base quality score recalibration and local realignment at known indels. The BWA aligner was used for mapping reads to the human genome build 38. Single nucleotide variants (SNVs) and insertions/deletions (indels) were jointly called across all samples using Genome Analysis Toolkit (GATK) HaplotypeCaller package version 4. Default filters were applied to SNV and indel calls using the GATK Variant Quality Score Recalibration (VQSR) approach. Annotation was performed using Variant Effect Predictor (VEP). The variant call sets were uploaded to seqr (https://seqr.broadinstitute.org/) for collaborative analysis between the Center for Mendelian Genomics (CMG) and investigator. Identified variants were validated with Sanger sequencing run in all available members of the family. For Case C, WES was performed in Mendelics Genomic Analysis facilities using Illumina NovaSeq 6000. Sequencing library was built with Illumina Nextera Flex and for capture of target regions Customized Exome Kit from Twist Biosciences was used. Sequencing of sample resulted in a total of 27.173.797 paired 101 bp sequences mapped to b37/hg19 human genome reference. 96.1 % of the Nextera exome reference was sampled at 10X or more. Exome reads were mapped to b37/hg19 reference using BWA MEM software (http://bio-bwa.sourceforge.net/). Resulting BAM files were genotyped using Broad Institute best practices with GATK (https://software.broadinstitute.org/gatk/). Resulting VCF files were processed using Mendelics in-house pipeline for annotation and filtering. "In silico" pathogenicity evaluation was performed using VSA, a Mendelics proprietary machine-learning based software. Aligned BAM files were also processed by ExomeDepth in order to identify CNVs. The heterozygous c.2771T>C; p.Leu24Pro variant in *ATP1A3* was validated in the proband by Sanger sequencing. The proband's parents were also investigated for the *ATP1A3* variant using Sanger sequencing, which was found to be absent in both, in peripheral blood samples. For Case D, WES was performed at GeneDx. Genomic DNA from the submitted specimen was enriched for the complete coding regions and splice site junctions for most genes of the human genome using a proprietary capture system developed by GeneDx for next-generation sequencing with CNV calling (NGS-CNV). The enriched targets were simultaneously sequenced with paired-end reads on an Illumina platform. Bi-directional sequence reads were assembled and aligned to reference sequences based on NCBI RefSeq transcripts and human genome build GRCh37/UCSC hg19. Using a custom-developed analysis tool (XomeAnalyzer), data were filtered and analyzed to identify sequence variants and most deletions and duplications involving three or more coding exons (Retterer et al., 2016).

**Phenotypic assessment**

All affected individuals and clinical data were examined by neurologists and/or geneticists, and radiologists, and polymicrogyria was diagnosed using criteria described previously (Jansen et al., 2015; Smith and Walsh, 2020). Supplemental text below summarizes their phenotypes and detailed clinical and radiographic evaluations.

**Human brain tissue preparation and mRNA *in situ* hybridization**

Performed as previously described Smith et al[3]. Briefly, following fixation (4% PFA) and cryoprotection (30% sucrose), brains were frozen using Isopentane on dry ice. Samples were sectioned at 20 – 30 µm thickness (Leica Cryostat), mounted immediately onto warm charged SuperFrost Plus slides (Fisher), and stored at –80°C. We followed manufacturer's standard protocol for multiplex fluorescent *in situ* hybridization (Multiplex Version 2 kit, Advanced Cell Diagnostics). *In situ* probe ACD catalog numbers are as follows: *ATP1A3*: 503941, *Vimentin*: 479411, *Eomes*: 429691. Whole tissue mRNA *in situ* imaging was performed on a Zeiss Axio Observer with automated image. (**Figure 3B**) Bright-field images were background corrected by Zen Blue Software for center intensity illumination and stitched together (**Figure 3B**).

**Bulk human cortex gene expression analysis**

The Allen Human Brain Atlas (ABA) publishes a rich dataset of cortical genetic expression across cortical brain regions, from age 8 weeks post conception to adult ages.(Jones et al., 2009) BrainSpan data analysis of *ATP1A1* (chr1:116,915,289-116,952,883, GRCh37/hg19), *ATP1A2* (chr1:160,085,548-

160,113,381, GRCh37/hg19), and *ATP1A3* (chr19:42,470,733-42,498,384, GRCh37/hg19) was performed. *ATP1A4* is not expressed in the CNS and was not included as the read counts were near zero. RNA-seq expression measured in RPKM (reads per kilobase exon per million mapped reads) was obtained from the BrainSpan project data and summarized to Gencode v10 exons for all annotated neocortical tissues aged 12 weeks post conception to 36 years. To obtain a moving average across ages, we fit a polynomial to the data using Igor pro software. Brain regions for Figure 3B include: dorsolateral prefrontal cortex; ventrolateral prefrontal cortex; anterior (rostral) cingulate (medial prefrontal) cortex; orbital frontal cortex; primary motor-sensory cortex; parietal neocortex; posterior (caudal) superior temporal cortex (area 22c), inferolateral temporal cortex (area 20); occipital neocortex, thalamic regions and hippocampus, among others.

**Single-cell suspensions for Drop-Seq**

The intact left brain hemisphere of a 21 wpc specimen was transported to the McCarroll lab in ice-cold Hibernate-E medium (ThermoFisher). The tissue was transferred to a dissection dish and submerged in ice-cold L-15 Medium (ThermoFisher) supplemented with 10% FBS (ThermoFisher). The medium was kept refrigerated throughout the dissection. Upon removal of the meninges, the hemisphere was divided along the rostro-caudal axis into 6 serial coronal slabs, each ~5cm thick, spanning the rostro-caudal axis. Regions of interest (Fig S2, Fig 4) were microdissected under visual guidance of a stereoscope (Leica MZ10), and identified using a fetal human atlas as reference.(Bayer et al., 2005) Single cell suspensions were produced using the MACS Neural Tissue Dissociation kit [papain (P), Miltenyi Biotec)] as described (Florio et al., 2015) with modifications. Briefly, tissue fragments containing regions of interest were transferred into 15 mL falcon tubes containing 2 mL of "Enzyme mix 1" (containing papain) and incubated at 37 C for 20' in a benchtop incubator without agitation. Tubes containing digested tissue were then transferred onto ice and 30 μL of "Enzyme mix 2" (containing papain inhibitor) were added to each sample. Tissue chunks were carefully titrated with a P1000 pipette, the triturated cells were centrifuged at 300 g for 5', and the cell pellets were resuspended in 1x PBS containing 0.01% BSA. Cell suspensions were then passed through a pre-wet 40 μm filter into a new tube on ice, and diluted to a final concentration of 220 cells/μl for Drop-Seq.

**Single-nucleus suspensions for Drop-Seq**

Nuclei were isolated from frozen cortical tissue (240-days-old donor), from 4 anatomically distinct cortical regions (BA10, BA41/42, BA17, BA40, Fig 5), dissected by NIH BioBank and stored in individual microcentrifuge tubes at -80°C. Single-nucleus suspensions and processed for DropSeq as described in detail at: https://protocols.io/view/extraction-of-nuclei-from-brain-tissue-2srged6, with modifications detailed (Krienen et al., 2019).

**Drop-seq sequencing**

Single-cell cDNA libraries were processed and sequenced using Illumina short read sequencing on a Nova-seq platform at the Broad Institute. Libraries were sequenced at an average depth of >8 reads per UMI. Sequencing reads were aligned to the hg19 human reference genome. For the fetal dataset, only reads mapping to exons were used; for the postnatal dataset, reads mapping to either exons and introns were used. Software and core computational analysis for alignment and downstream processing of Drop-seq sequencing reads are freely available at https://github.com/broadinstitute/Drop-seq/releases.

**Independent component analysis and clustering**

Cell selection was performed as described (Saunders et al., 2018) for each sequencing pool independently and the resulting digital gene expression (DGE) matrices were then merged into unique "dataset DGEs" (i.e. fetal and postnatal) which were analyzed separately. Independent component analysis (ICA) was performed using the R' fastICA package on each dataset's DGE, after normalization and variable gene selection, as described in Saunders 2015 and Krienen 2019, with modifications. Single-cell/nucleus libraries containing fewer than 500 genes were excluded from the analysis, and genes encoding Mitochondrial RNAs ('^RPS'), Ribosomal proteins ('^RPS', '^RPL'), and Histones ('^HIST) were filtered out from the DGEs prior to ICA. The number of significant components used for clustering was determined using the JackStraw function from Seurat (https://www.rdocumentation.org/packages/Seurat/versions/3.1.4). Cells identified as doublets or outliers were iteratively removed through multiple rounds of clustering. Clusters were identified using canonical cell type markers (Fig S2, S4), and only neural clusters (i.e. excitatory neurons, inhibitory neurons, glia, and neural progenitor cells) were selected in the analysis. A final round of clustering was then run on the filtered DGEs, and the optimal clustering resolutions were chosen based on the salience of unique markers within each cluster, as described (Saunders et al., 2018).

**Statistical analysis**

Data analysis was performed using R studio. Source code available by request. Gene expression normalization and scaling methods for each analysis are specified in figure legends. Clustering Heatmaps were generated using R's *pheatmap* package, using the 'ward.D2' clustering method and *euclidean* clustering distance. Pairwise gene correlation coefficients were calculated upon relative-count normalization and scaling using R's *cor* function. tSNE embeddings were generated as described in Saunders 2015. UMAP embeddings were generated using R's *uwot* package using as input a matrix wherein each row is a cell in the DGE and each column is the aggregated expression, for that given cell, of the genes positively loading onto each individual component used in ICA. The resulting matrixes was then scaled by cell using R'*scale* function. The UMAP algorithm was run with default parameters, with a n_neighbors equal to the square root of number of cells in the dataset. Pseudotime analysis was performed using *Monocle 3* (https://cole-trapnell-lab.github.io/monocle3/). The analysis was run using default parameters and using as input a subset of the fetal dataset's DGE containing only cells assigned to EN clusters.

**Graphics**

Graphics prepared in Adobe Illustrator and using BioRender software.

### SUPPLEMENTAL RESULTS

**Supplemental Text**

Clinical descriptions of Case A-D; Related to Figure 1

**Table S1**

PMG-associated *ATP1A3* alleles

**Figure S1**

Sequencing and conservation of Cases A-D with *ATP1A3* pathogenic variants; Related to Figure 1

**Figure S2**

Anatomical origin and gene-expression markers for cortical fetal samples used in DropSeq experiments; Related to Figure 4

**Figure S3**

Cell-type specific expression and pair-wise correlation of *ATP1A1-3* in the fetal human neocortex; Related to Figure 4

**Figure S4**

Clustering analysis of *ATP1A3* expression in the infant human cortex, Related to Figure 5

**Figure S5**

Pair-wise correlation of *ATP1A1-3* in the infant human neocortex; Related to Figure 5

**Table S2.**

Gene ontology analysis of the top-100 genes correlating with *ATP1A3* in the EN cluster in the postnatal cortex, Related to Figure 5

**Figure S6**

Comparison of *ATP1A3* expression within cortical interneurons isolated from mice, macaque, and human; Related to Figure 5

**Figure S7**

Schematic of *ATP1A3* variants with respect to disease severity

**Table S3**

Summary of published *ATP1A3* variants associated with disease with original references and phenotypic features

### CLINICAL DESCRIPTIONS

#### Case A: Bilateral Frontoparietal PMG (p.Arg901Met)

Case A, a female child of nonconsanguineous parents from the Azores of Portugal, was born at full term by spontaneous vaginal delivery following an unremarkable pregnancy. She required oxygen for respiratory distress at birth, exhibited neonatal jaundice and was discharged from intensive care after 10 days. Her developmental delays were noted at an early age, as she could not hold a pacifier. At 2 and 3 years of age, she had surgical corrections for strabismus. Drooling was a significant concern by 6 years of age. She had Botox injections in childhood for her spastic quadriparesis. Episodes suspicious for seizure activity occurred around 8.5 years and at about 9 years she developed focal impaired awareness seizures, which was treated with oxcarbazepine. Evaluation at 9 years and 2 months of age revealed a head circumference of 51 cm (-0.73 standard deviations, SD), length of 110.6 cm (-3.82 SD) and weight of 17.7 kg (-2.92 SD). She was developmentally and cognitively delayed, able to sit alone and walk using a walker with assistance, point and use some sign language but had no speech, and could hold a pencil but not write. Drooling was still an issue and she did not chew. A prior barium swallow study was negative for aspiration. She attended a public school in a contained class and received physical, occupational and speech therapies. Asymmetric spastic quadriparesis (left more significant than right) with hypertonia and athetoid movements of the hands and fingers were noted on examination, as were a high palate, thin upper lip and smooth philtrum.

Investigations included an EEG at 8 years 8 months of age that was abnormal and showed no clear anterior to posterior gradient. Alpha activity was more prominent over the frontal area with some attenuation of the amplitude and slightly faster frequencies over the posterior regions. A clear beta activity could not be identified over the anterior or central areas in a consistent manner. Throughout the recording, spikes and high amplitude sharp and slow waves were noted bilaterally over the frontotemporal areas, with the right being greater than left. Brain MRI performed at 5 months showed polymicrogyria (PMG) in the perisylvian cortex bilaterally, as well as extensive thinning of the spinal cord from T1-T12. MRI at 9 years and 3 months of age revealed PMG in the perisylvian cortex, which was thicker than at 5 months, as well as in the suprasylvian cortex, predominantly frontally but extending much farther rostrally. The parietal lobes were very abnormal and pachygyric in appearance. The posterior part of frontal and temporal cortex was abnormally thick, with somewhat shallow sulci, giving a pachygyric appearance. These abnormalities were bilateral but asymmetric, with the left more affected than the right. The most anterior aspects of the frontal and parietal lobes, as well as the medial and inferior cerebral cortex, were spared. However, it is worth noting the pachygyric appearance may be the result of averaging of a large segment of the cortex as a result of thick sections on a 20 year old MRI. The corpus callosum, basal ganglia and midline structures appeared normal. The cerebral white matter volume was significantly reduced compared to the 5-month study with otherwise normal myelination. Lateral ventricles were mildly enlarged, and the cerebellum was disproportionately large when compared to the cerebrum. The spinal cord was not included in the MRI at 9 years. Unremarkable genetic testing included a karyotype (46, XX), array CGH, and Fragile X, as well as *ARX* gene sequencing.

**Case B: Extensive Bilateral PMG (c.2921+1G>A)**

Case B, a male child of nonconsanguineous parents from the Philippines, was born at 38 weeks' gestation by Cesarean section due to decreased fetal heart rate following a pregnancy complicated by maternal gestational diabetes. At birth, his head circumference was 34 cm (-0.83 SD) and weight was 2.995 kg (-0.91 SD), and he developed seizures 12 hours after delivery. Extensive bilateral polymicrogyria was identified by brain MRI at 6 weeks of age and video EEGs confirmed a severe epileptic encephalopathy that proved resistant to essentially all antiepileptic drugs. EEGs at 2 months of age showed abundant electroclinical and electrographic seizures beginning predominantly from the right parasagittal region but also from the left and bilateral parasagittal regions. Some seizures had diffuse onset associated with head jerks, and interictally, there were independent epileptogenic discharges from the right and left parasagittal regions activated during sleep. Visual and auditory evoked potential studies were normal at 4 and 5 months of age respectively. He required nasogastric tube followed later by a G-tube for feeding.

Evaluation at 14 months revealed microcephaly, with a head circumference of 41.5cm (-4.63 SD); his weight was 10.4 kg (-1.61 SD) and height was 73cm (-0.38 SD). He exhibited severe global developmental delay (never rolled over, sat up or talked), intermittent nystagmus, significant axial and appendicular hypotonia, and hyporeflexia. His severe epileptic encephalopathy at 14 months was described by 24-hour video-EEG study as having a poorly organized and slow background with excessively frequent multifocal independent epileptiform discharges throughout almost all brain regions, though no electrographic seizures were noted on that study. Brain MRI at 14 months noted widespread bilateral polymicrogyria with some calcification at the cortical-white matter junction. While myelination was slightly delayed at 2 months, it was markedly delayed at 14 months with progressive volume loss, and the corpus callosum was normally formed but thin. The basal ganglia appeared atrophic compared to the initial scan. Initial MRI studies revealed a small, cyst-like (perhaps cavitation) areas observed within the hippocampal heads and bodies (left side slightly larger). Follow-up studies the following year demonstrate hippocampi are much smaller with what appears to be central atrophy (possibly shrinkage after cavitation) of the hippocampal head and ventral body. Adjacent white matter also shrinking but, most likely, gray matter (hippocampi) shrunk first. Other examinations included a karyotype and 22q11 FISH assay, chromosomal microarray, *ADGRG1* (*GPR56*) gene sequencing, urine organic and plasma amino acid testing, acylcarnitine profile and very long chain fatty acid testing, which were all unremarkable.

**Case C: Unilateral PMG (p.Leu924Pro)**

Case C is a male born to nonconsanguineous parents from Brazil. His 32-year-old mother has a personal history of hypothyroidism. After an uneventful pregnancy, he was born at 38 weeks of gestation by Cesarean section for acute fetal distress, weighed 3.66 kg (0.2 SD) and had Apgar scores of 6 and 8 at 1 and 5 minutes, respectively. Hypotonia and a left clubfoot (for which casting was indicated) were noted at birth. Episodes of upward rolling of the eyes on the first day of life prompted admission to the NICU. An EEG confirmed the presence of electroclinical seizures and phenobarbital was initiated. On day 3, he exhibited apnea and hypertonia, however 24-hour video-EEG study failed to detect any significant abnormalities. Neurological exam revealed episodic dystonic posturing of the upper left extremity, alternating with excessive movements of closing his left hand with thumb adduction. Three electroencephalographic focal seizures and a clinical epileptic episode characterized by eye blinking and apnea occurred on day 4, and an EEG on day 5 described the epileptic seizures as periodic discharges with right posterior projection. Brain MRI, also on day 5, showed extensive right hemispheric polymicrogyria involving the right frontal, parietal, and temporal lobes, and the insula, as well as a small foci of signal abnormality in the periventricular white matter, compatible with neonatal ischemic injury. He was discharged at 13 days of age with a phenobarbital and levetiracetam regime.

His subsequent neuropsychomotor development was delayed and he required hospital admission for epileptic episodes at 2 and 4 months of age. Two EEGs around 4 months showed: (1) moderate periodic epileptic activity localized to the left temporal region and more rarely bilateral central projection, and low delta wave surges with left hemisphere projection (occasionally rhythmically occurring) suggestive of a localized rhythmic delta activity pattern; and (2) infrequent slow wave surges bilaterally in the anterior regions, and epileptic paroxysms with sharp wave morphology in the left temporal region and rare occurrences in the posterior regions of the left hemisphere and midline region. He was admitted again at 5 months due to bradycardia, without associated seizures, ultimately went into cardiogenic shock due to acute viral myocarditis and was discharged after hemodynamic recovery. Upon evaluation at 8 months of age, he was on a ketogenic diet and in use of Cannabidiol 4.4 mg/kg/day, Topiramate 50 50 16.5 mg/kg/day, phenobarbital 7.25 mg/kg/day. His examination revealed a head circumference of 42.5 cm (-1.75 SD), global hypotonia with present reflexes, very poor eye contact, babbling but no head control, rolling or gripping of objects.

**Case D: Extensive Multifocal Bilateral PMG (p.Gln851Arg)**

Case D is a male born to nonconsanguineous parents of European decent by emergency Cesarean section at 41 weeks gestation following a pregnancy complicated by fetal hydronephrosis. He had Apgar scores of 1 and 8 at 1 and 5 minutes respectively, and a birth weight of 3.8 kg (0.49 SD). He required a 30-day neonatal hospital stay, as his early course was complicated by seizures and he was diagnosed with a neonatal encephalopathy requiring cooling. Brain MRI soon after birth revealed bilateral polymicrogyria involving the right cerebral hemisphere more extensively than the left, most severe in the right fronto-temporo-parietal lobes and insula with relative sparing of the right occipital lobe. There was progression of diffuse white matter volume loss in the right cerebral hemisphere, right basal ganglia and corresponding progressive volume loss in the right cerebral peduncle, right aspect of the pons and upper medulla. Progression of *ex vacuo* dilatation of the right lateral ventricle was also noted, as was a small corpus callosum, especially the splenium. There was mild scattered ethmoid sinus mucosal thickening, and no evidence of an intracranial mass, acute intracranial hemorrhage or infarct, or extra axial fluid collection.

At 4 months of age, he underwent surgery for congenital hip dysplasia. At that time, he was noted to have frequent breath-holding spells, which worsened after the surgery. A video EEG then showed slowing during these spells and they were initially not considered to be epileptiform. His seizures are characterized by back and forth eye movements and sometimes involve shaking of his hands. He is treated with Phenobarbital, Keppra and Trileptal, is reported to have persistent intermittent abnormal eye movements and has spastic left hemiparesis, mainly involving his left arm but also his face.

At 15 and 30 months of age, head circumference was 42.7 cm (-3.5 SD) and 44.2 cm (-3.3 SD), respectively. On examination at 18 months of age, he exhibited very low tone with poor head control. Developmentally, he babbled, could not roll over, sat with support and tended to play with his right hand. He had some persistent feeding problems, was not yet receiving any solid food and was reported to have constant constipation. He was described to sleep like a newborn, waking up during the night and asking to be fed. He receives speech, physical and occupational therapies.

**Table S1, PMG-associated *ATP1A3* alleles**

| | Case A | Case B | Case C | Case D |
|---|---|---|---|---|
| **Genomic coordinate (GRCh38)** | **g.41968902C>A** | **g.41967661C>T** | **g.41968833A>G** | **g.41969571T>C** |
| **cDNA notation (NM_152296.4)** | **c.2702G>T** | **c.2921+1G>A** | **c.2771T>C** | **c.2552A>G** |
| **Protein notation** | **p.Arg901Met** | **NA** | **p.Leu924Pro** | **p.Gln851Arg** |
| **PhyloP** | **7.76 [-20.0;10.0]** | **NA** | **9.29 [-20.0;10.0]** | **9.29 [-20.0;10.0]** |
| **SIFT (v6.2.0)** | **Deleterious (score: 0, median: 3.46)** | **NA** | **Deleterious (score: 0.01, median: 3.46)** | **Deleterious (score: 0.01, median: 3.45)** |
| **Mutation Taster (v2013)** | **Disease causing (prob: 1)** | **NA** | **Disease causing (prob: 1)** | **Disease causing (prob: 1)** |
| **PolyPhen-2** | **Probably damaging (1.000)** | **NA** | **Probably damaging (1.000)** | **Probably damaging (1.000)** |
| **Splice predictions at nearest natural splice junction** | **NA** | **Predicted change at donor site 1 bps upstream: -100.0%MaxEnt: -100.0% NNSPLICE: -100.0% SSF: -100.0%** | **NA** | **NA** |
| **gnomAD v.2.1.1 (MAF)** | **0** | **0** | **0** | **0** |

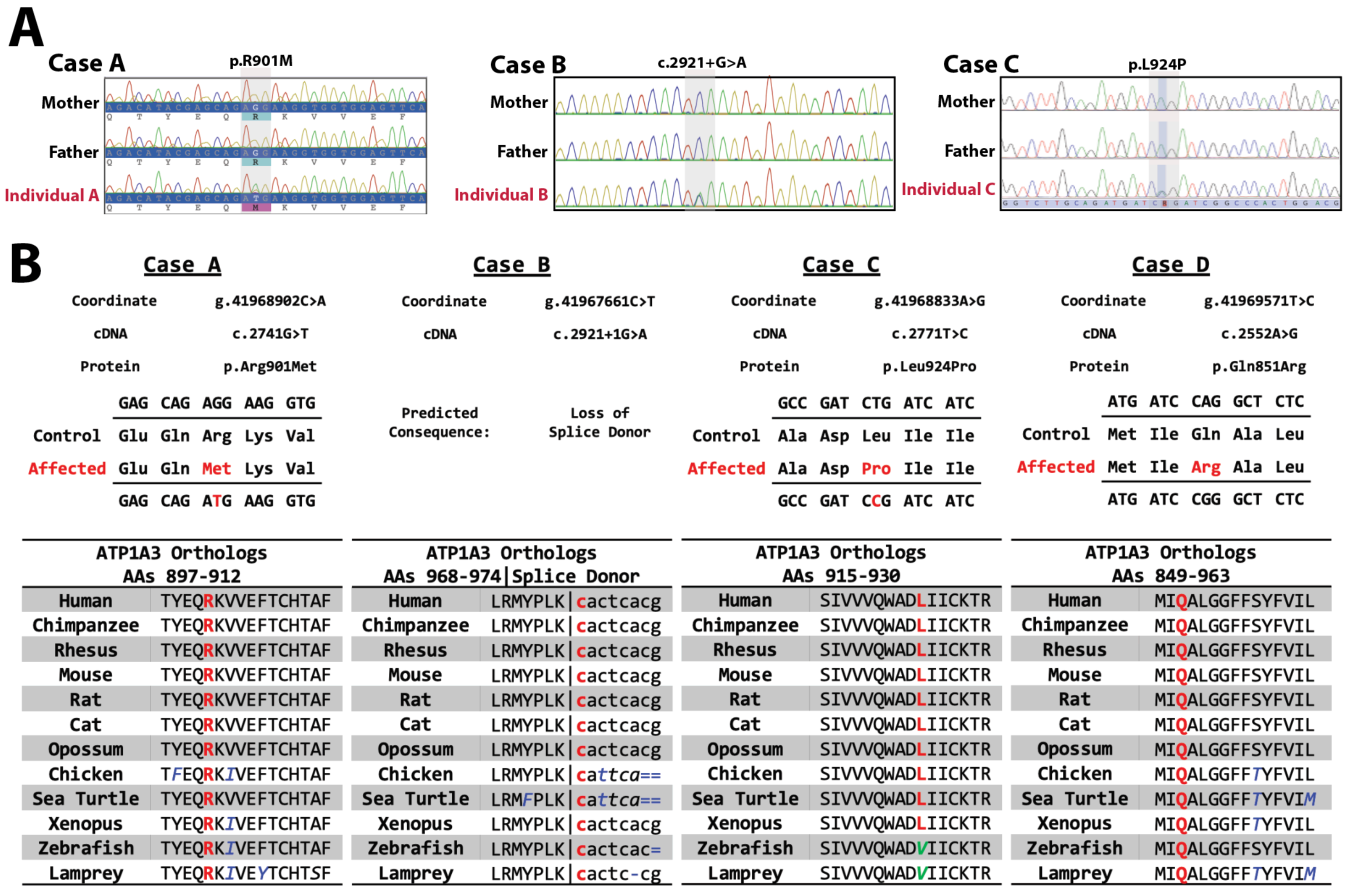


**Figure S1. Sequencing and conservation of Cases A-D with *ATP1A3* pathogenic variants; Related to Figure 1**

**Figure S1. Sequencing and conservation of Cases A-D with *ATP1A3* pathogenic variants; Related to Figure 1**

**(A)** *Left*, Representative *ATP1A3* Sanger chromatograms from unaffected (black text) parents and affected individual A, possessing a *de novo* heterozygous G>T substitution. . *Center*, Case B Sanger chromatograms from unaffected (black text) parents and affected *de novo* individual B, possessing a *de novo* heterozygous G>A substitution. *Right*, Case C Sanger chromatograms from unaffected (black text) parents and affected *de novo* individual C, possessing a *de novo* heterozygous T>C substitution. **(B)** *ATP1A3* sequence alignments of the amino acids surrounding the PMG associated mutations from Na,K-ATPase alpha-3 isoform ortholog across species, showing high degree of conservation.

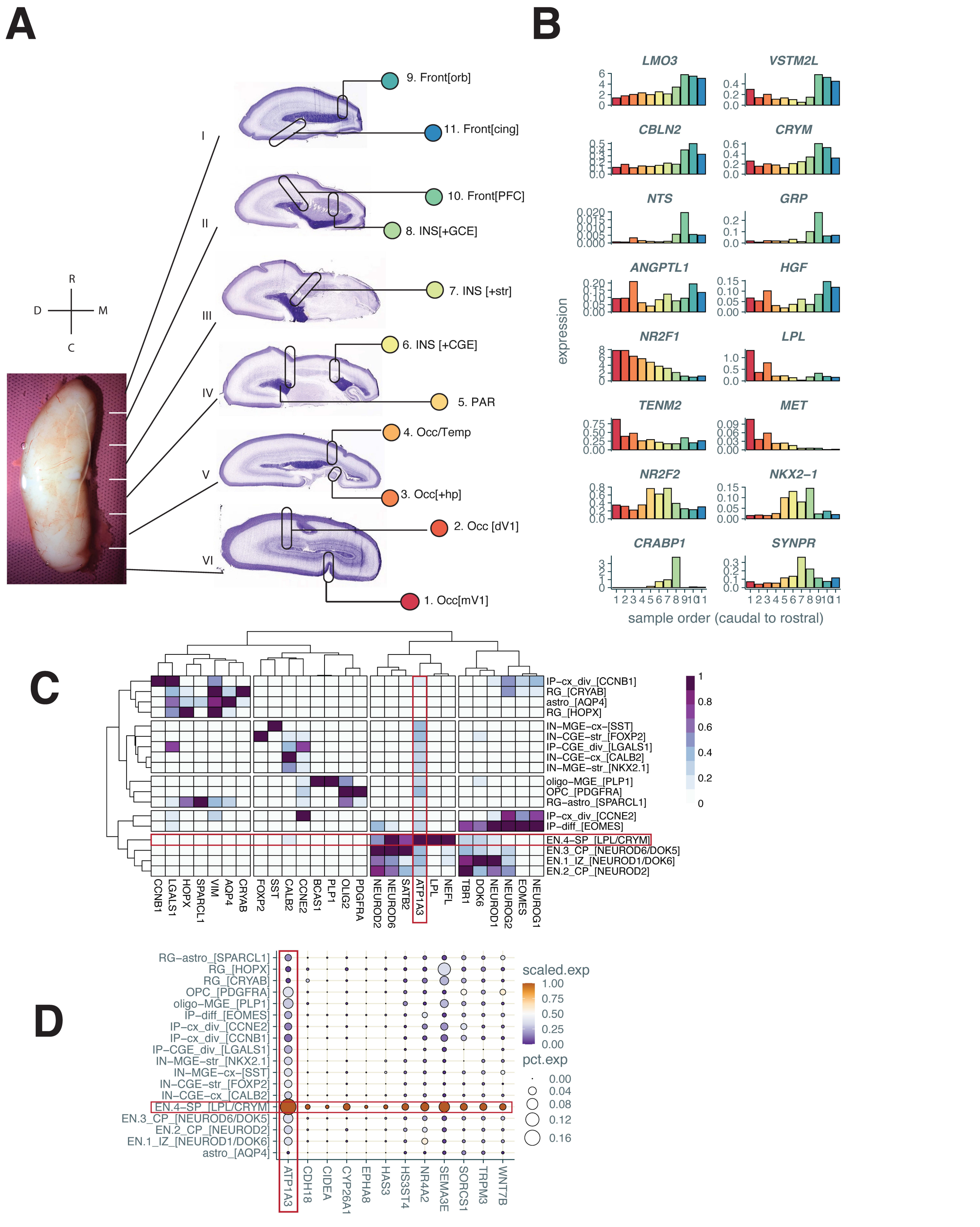


**Figure S2, Anatomical origin and gene-expression markers for cortical fetal samples used in DropSeq experiments; Related to Figure 4**

**Figure S2, Anatomical origin and gene-expression markers for cortical fetal samples used in DropSeq experiments;** Related to Figure 4

(**A**) *Left*, Photograph of the fetal neocortical specimen (21 wpc) used for DropSeq, prior to microdissection. Anatomical axes are indicated (R: rostral, C: caudal, D: dorsal, M: medial). Horizontal white lines indicate approximate location of dissection boundaries between cortical slabs (I-VI). *Right*, Representative Nissl-stained coronal section images of a 21wpc neocortex depicting anatomical detail and location of samples [source: Atlas of the Developing Human Brain, *BrainSpan* (www.brainspan.org). Fr: Frontal; Par: Parietal; Temp: Temporal; Occ: Occipital; Orb: Orbital Frontal Cortex; Ins: Insula; Cing: Cingulate cortex; dV1 and mV1: dorsal and medial Primary Visual Cortex, respectively; Str: Striatum; Hp: Hippocampus; CGE and MGE: caudal and medial Ganglionic Eminence, respectively. ***(B)*** Bar graphs showing relative expression of regional marker genes (y-axis) across samples (x-axis, samples 1-11, ordered from caudal to rostral). Note the enrichment *of LMO3, VSTM2L, CBLN2, CRYM, NTS, GRP* in frontal and prefrontal cortex samples (samples 9-11); *NR2F1, LPL, TENM2, MET* in occipital cortex samples; *ANGPTL1, HGF* in medial occipital cortex samples including portion of hippocampus; *NR2F2* in temporo-parietal cortex samples; *NKX2-1* in medial temporal cortex samples including MGE; *CRABP1* in medial cortex samples including CGE; and *SYNPR* in insular cortex samples including striatum. Expression of each gene was sum-aggregated by sample, then gene expression was relative-count normalized (divided by total counts/sample x 100k). **(C)** Heatmap showing hierarchical clustering and expression specificity of selected cell-type markers (columns) in the different clusters (rows). Note the highest relative expression of *ATP1A3* is the EN.4 SP cluster, marked by deep-layer EN marker *NEFL* and SP marker LPL. Expression of each gene was sum-aggregated by cluster, relative-count normalized, then rescaled from 0 to 1. **(D)** Dot-plot showing specific enrichment of subplate markers in the EN.4 SP cluster. Color scale codes for mean gene expression by cluster; size of the dots codes for percentage of cells expressing a given gene in each cluster. RG: radial glia IP: intermediate progenitors (div: dividing, diff: differentiating, cx: cortex, str: striatum); OPC: oligodendrocyte progenitor cell; IN: inhibitory neurons, EN: excitatory neurons (CP: cortical plate, IZ: intermediate zone); astro: astrocytes; OPC: oligodendrocyte progenitor cells.

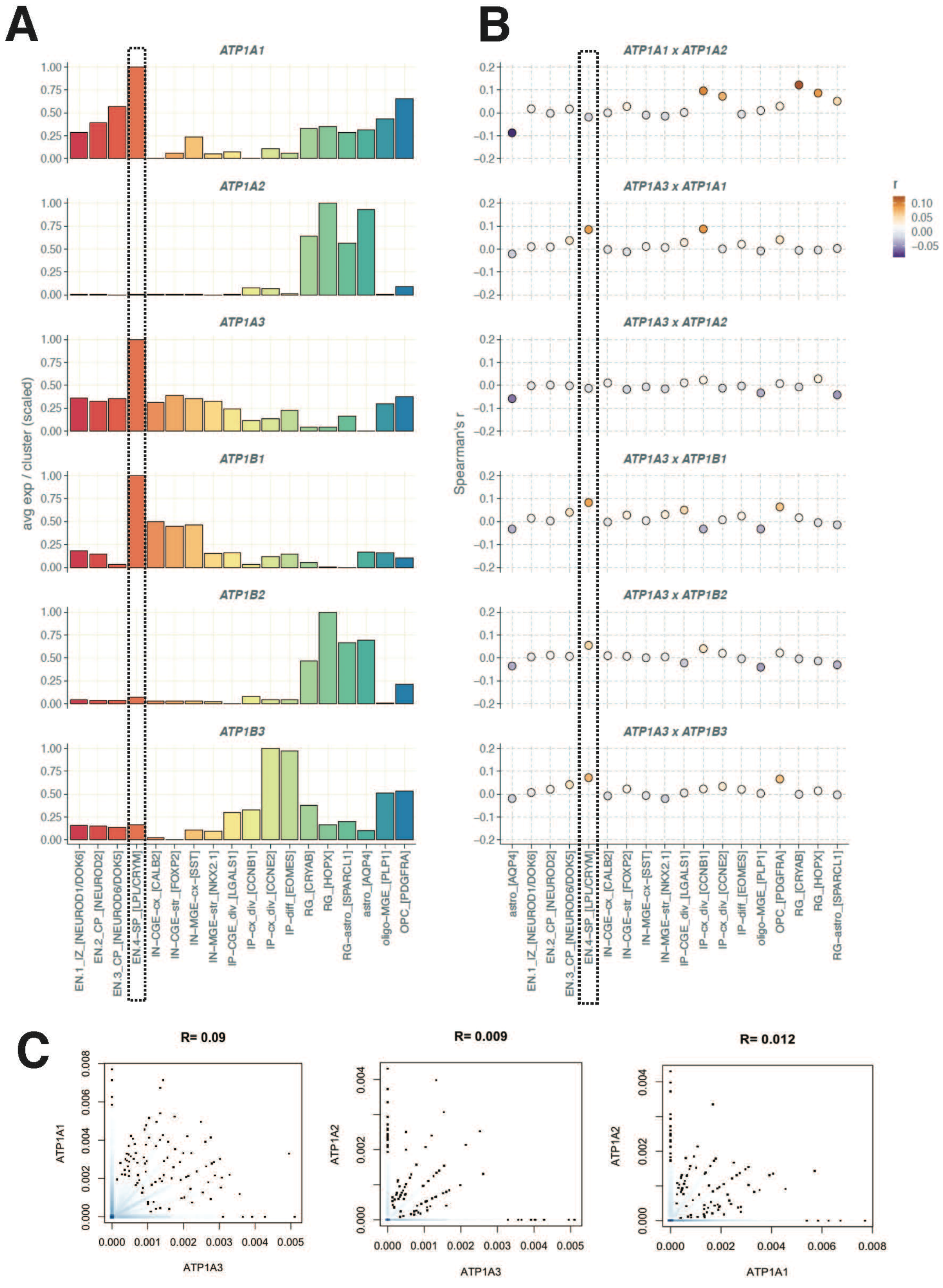


**Figure S3 Cell-type specific expression and pair-wise correlation of *ATP1A1-3* in the fetal human neocortex; Related to Figure 4**

**Figure S3 Cell-type specific expression and pair-wise correlation of *ATP1A1-3* in the fetal human neocortex; Related to Figure 4**

(**A)** Bar graphs showing relative expression of *ATP1A1*, *ATP1A2* and *ATP1A3* (y-axis) across clusters. Gene expression (y-axis) was normalized by cell (divided by total UMIs/cell x 100k), then the mean value for each gene was aggregated by cluster and rescaled from 0 to 1. Note the enrichment of *ATP1A1* and *ATP1A3* in the EN4.SP cluster, and the enrichment of *ATP1A2* in glial clusters (anti-correlating with *ATP1A3*). **(B)** Dot plots showing pairwise correlation coefficients between *ATP1A1, ATP1A2* and *ATP1A3* expression across clusters. Dots represent Spearmann's correlation coefficients (r), color coded by association strength. Note the positive correlation between expression of *ATP1A3* and *ATP1A1*, but not *ATP1A2*, in the EN SP cluster. **(C)** Scatter plots showing pairwise comparisons of *ATP1A1*, *ATP1A2* and *ATP1A3* expression in single cells. Each dot represents a cell. Gene expression was normalized by cell. Pearson correlation coefficients (R) for each pair are indicated above each plot.

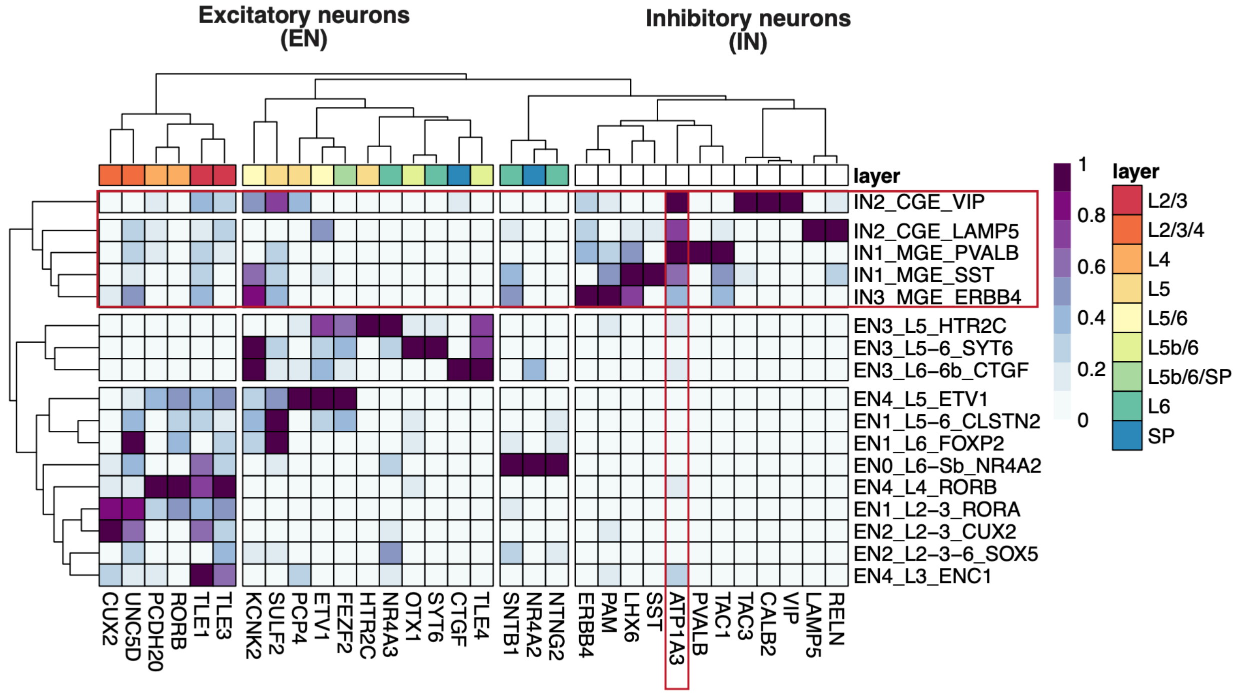


**Figure S4. Clustering analysis of *ATP1A3* expression in the postnatal cortex, Related to Figure 5**

**Figure S4. Clustering analysis of *ATP1A3* expression in the postnatal cortex, Related to Figure 5**

8-month old neocortex of 51,878 single nuclei profiled by DropSeq. EN: excitatory neurons; IN: inhibitory neurons. Heatmap showing hierarchical clustering and expression specificity of selected cell-type markers (columns) in the different clusters (rows). Mean gene expression was aggregated by cluster, then rescaled from 0 to 1. Representative cortical marker genes including EN layer 2-3 marker CUX2, EN layer-5 marker FEZF2, EN layer-4 marker RORB, MGE-derived IN markers SST+, VIP+, CGE-derived IN maker LAMP5.

| Enrichment FDR | Genes in list | Total genes | Functional Category | Genes |
|---|---|---|---|---|
| 3.17856230303671E-25 | 59 | 1808 | Neuron part | STMN4 GABRA1 SYT1 HSP90AA1 PEBP1 SLC32A1 STMN2 CPE HSPA8 GABRG2 EPHA4 STMN1 MAP1B SYT11 RAC1 GPM6A THY1 SLC6A1 SV2A NPTXR CD200 CLU APP SERPINI1 PRNP GRIN1 DNER ATP2B1 [illegible] |
| 1.02200062088446E-21 | 47 | 1268 | Synapse | ATP2B1 RAC1 YWHAZ NPTXR GABRA1 SYT1 PEBP1 SLC32A1 CPE HSPA8 GABRG2 EPHA4 MAP1B SV2A CLSTN1 APP MEF2C SCAMP1 RABAC1 NRXN2 RTN4 CLU CANX YWHAH OLFM1 NDFIP1 SYT11 RTN3 [illegible] |
| 5.4070009738657E-21 | 80 | 4252 | Vesicle | MDH1 PKM ATP2B1 ATP6AP1 HSP90AA1 APLP2 DNAJA1 FTL GNAS PEBP1 HSP90AB1 PLD3 ALDOC CPE HSPA8 GAPDH LDHB STMN1 CLU SEPT7 CANX YWHAH ARF3 NARS ITM2B RAC1 TUBB2A APP ATP1B1 [illegible] |
| 1.5301082324951E-20 | 39 | 879 | Somatodendritic compartment | GABRA1 HSP90AA1 PEBP1 SLC32A1 CPE HSPA8 GABRG2 EPHA4 MAP1B RAC1 GPM6A THY1 NPTXR CD200 STMN2 CLU APP SERPINI1 PRNP GRIN1 DNER ATP2B1 GNAS HSP90AB1 RTN4 CANX OLFM1 NDFIP1 [illegible] |
| 4.56114050184702E-18 | 44 | 1371 | Neuron projection | STMN4 GABRA1 SYT1 PEBP1 SLC32A1 STMN2 HSPA8 GABRG2 EPHA4 STMN1 MAP1B SYT11 RAC1 GPM6A THY1 SLC6A1 SV2A NPTXR CD200 CLU APP PRNP GRIN1 DNER ATP2B1 HSP90AA1 GNAS HSP90AB1 [illegible] |
| 8.39569886146249E-18 | 59 | 2625 | Cytoplasmic vesicle | SYT1 SCAMP1 CHGB PEBP1 SLC32A1 CPE SYT11 RAC1 SV2A NSG2 SCG2 CLIP3 ITM2B APP SERPINI1 TUBA1A DNER PSAP VAMP2 GABRA1 GABARAPL2 HSP90AA1 GNAS HSP90AB1 SCG3 STMN2 RABAC1 [illegible] |
| 8.39569886146249E-18 | 59 | 2628 | Intracellular vesicle | SYT1 SCAMP1 CHGB PEBP1 SLC32A1 CPE SYT11 RAC1 SV2A NSG2 SCG2 CLIP3 ITM2B APP SERPINI1 TUBA1A DNER PSAP VAMP2 GABRA1 GABARAPL2 HSP90AA1 GNAS HSP90AB1 SCG3 STMN2 RABAC1 [illegible] |
| 5.59716225012485E-17 | 37 | 1004 | Synapse part | RAC1 GABRA1 SYT1 PEBP1 SLC32A1 CPE HSPA8 GABRG2 EPHA4 SV2A CLSTN1 APP ATP2B1 MEF2C SCAMP1 RABAC1 NRXN2 RTN4 CANX YWHAH MAP1B SYT11 FXYD6 ARF1 GPM6A SLC6A1 CALM3 CADM3 [illegible] |
| 2.4595883190098E-16 | 79 | 4988 | Endomembrane system | RTN4 SEC62 SYT1 ERLEC1 SCAMP1 CHGB PEBP1 SLC32A1 CPE TMEM30A NDFIP1 SYT11 ARF3 ITM2B ARF1 SV2A HSP90B1 SOAT2 NSG2 ARL6IP1 SCG2 GABARAPL2 GNAS PTHLH CLIP3 GAPDH CLU CANX [illegible] |
| 2.70863924797889E-16 | 29 | 594 | Cell body | HSP90AA1 PEBP1 CPE MAP1B GPM6A NPTXR CD200 STMN2 SERPINI1 TUBB ATP2B1 HSP90AB1 RTN4 EPHA4 CANX OLFM1 SYT11 APP CRH THY1 UCHL1 SV2A CKB ENC1 SEZ6L2 CDK5R1 ATP6AP2 DNER [illegible] |
| 1.46533344571598E-15 | 52 | 2300 | Extracellular exosome | MDH1 PKM ATP2B1 ATP6AP1 HSP90AA1 APLP2 DNAJA1 FTL GNAS PEBP1 HSP90AB1 PLD3 ALDOC CPE HSPA8 GAPDH LDHB STMN1 CLU SEPT7 CANX YWHAH ARF3 NARS ITM2B RAC1 TUBB2A APP ATP1B1 [illegible] |
| 1.99712446442791E-15 | 52 | 2326 | Extracellular organelle | MDH1 PKM ATP2B1 ATP6AP1 HSP90AA1 APLP2 DNAJA1 FTL GNAS PEBP1 HSP90AB1 PLD3 ALDOC CPE HSPA8 GAPDH LDHB STMN1 CLU SEPT7 CANX YWHAH ARF3 NARS ITM2B RAC1 TUBB2A APP ATP1B1 [illegible] |
| 1.99712446442791E-15 | 52 | 2324 | Extracellular vesicle | MDH1 PKM ATP2B1 ATP6AP1 HSP90AA1 APLP2 DNAJA1 FTL GNAS PEBP1 HSP90AB1 PLD3 ALDOC CPE HSPA8 GAPDH LDHB STMN1 CLU SEPT7 CANX YWHAH ARF3 NARS ITM2B RAC1 TUBB2A APP ATP1B1 [illegible] |
| 4.60386356513867E-15 | 26 | 512 | Presynapse | SYT1 PEBP1 SLC32A1 HSPA8 SV2A ATP2B1 SCAMP1 RABAC1 NRXN2 EPHA4 CANX YWHAH SYT11 FXYD6 APP GPM6A SLC6A1 CALM3 CADM3 YWHAG CDK5R1 GRIN1 ACTG1 CALM1 VDAC1 VAMP2 |
| 5.08793535533338E-15 | 26 | 517 | Neuronal cell body | HSP90AA1 PEBP1 CPE MAP1B GPM6A NPTXR CD200 STMN2 SERPINI1 ATP2B1 HSP90AB1 RTN4 EPHA4 CANX OLFM1 APP CRH THY1 UCHL1 SV2A CKB ENC1 SEZ6L2 CDK5R1 DNER PGRMC1 |
| 5.08793535533338E-15 | 36 | 1108 | Secretory vesicle | SYT1 CHGB PEBP1 SLC32A1 CPE SYT11 SV2A SCG2 SCAMP1 SCG3 RABAC1 CLU APP CALM3 SERPINI1 SCG5 GRIN1 CALM1 VDAC1 VAMP2 PKM HSP90AA1 APLP2 FTL HSP90AB1 PGRMC1 ALDOC HSPA8 [illegible] |
| 5.81546312965533E-14 | 20 | 293 | Distal axon | PEBP1 STMN2 HSPA8 GPM6A NPTXR STMN4 HSP90AA1 HSP90AB1 SLC32A1 EPHA4 MAP1B SYT11 APP THY1 CALM3 CDK5R1 ACTG1 CALM1 OLFM1 GRIN1 |
| 7.76643366991935E-14 | 27 | 639 | Axon | SYT1 PEBP1 STMN2 HSPA8 SYT11 GPM6A SLC6A1 NPTXR CD200 STMN4 HSP90AA1 HSP90AB1 SLC32A1 GABRG2 EPHA4 CANX OLFM1 MAP1B APP CRH THY1 UCHL1 CALM3 CDK5R1 ACTG1 CALM1 GRIN1 |
| 7.76643366991935E-14 | 45 | 1930 | Whole membrane | GABARAPL2 SYT1 SCAMP1 PEBP1 SLC32A1 THY1 SV2A TOMM20 VDAC1 CLIP3 LDHB ITM2B RAC1 APP PRNP SLC25A5 ATP2B1 SLC22A17 HSP90AB1 SCG3 CPE TMEM30A EPHA4 ATP6V0B TUBA1B NDFIP1 [illegible] |
| 7.79073549376435E-14 | 49 | 2287 | Cell projection | STMN4 GABRA1 SYT1 PEBP1 SLC32A1 STMN2 HSPA8 GABRG2 EPHA4 STMN1 MAP1B SYT11 RAC1 GPM6A THY1 SLC6A1 SV2A NPTXR CD200 CLU SEPT7 APP PRNP GRIN1 DNER THSD7A PKM ATP2B1 HSP90AA1 [illegible] |
| 9.35400158788307E-14 | 27 | 647 | Dendrite | GABRA1 SLC32A1 HSPA8 GABRG2 EPHA4 MAP1B RAC1 THY1 NPTXR CLU APP PRNP GRIN1 DNER ATP2B1 HSP90AA1 GNAS HSP90AB1 CANX NDFIP1 SYT11 GPM6A SV2A CKB NSG2 CLSTN1 CDK5R1 |
| 9.35400158788307E-14 | 22 | 392 | Axon part | PEBP1 STMN2 HSPA8 GPM6A NPTXR STMN4 HSP90AA1 HSP90AB1 SLC32A1 EPHA4 MAP1B SYT11 APP CRH THY1 UCHL1 CALM3 CDK5R1 ACTG1 CALM1 OLFM1 GRIN1 |
| 9.35400158788307E-14 | 27 | 649 | Dendritic tree | GABRA1 SLC32A1 HSPA8 GABRG2 EPHA4 MAP1B RAC1 THY1 NPTXR CLU APP PRNP GRIN1 DNER ATP2B1 HSP90AA1 GNAS HSP90AB1 CANX NDFIP1 SYT11 GPM6A SV2A CKB NSG2 CLSTN1 CDK5R1 |
| 3.34448717619936E-13 | 70 | 4617 | Extracellular region | MDH1 PKM ATP2B1 ATP6AP1 HSP90AA1 APLP2 DNAJA1 FTL GNAS PEBP1 HSP90AB1 PLD3 ALDOC CPE HSPA8 GAPDH LDHB STMN1 CLU SEPT7 CANX YWHAH ARF3 NARS ITM2B RAC1 TUBB2A APP ATP1B1 [illegible] |
| 3.34448717619936E-13 | 39 | 1523 | Cell projection part | GABRA1 PEBP1 SLC32A1 STMN2 HSPA8 GABRG2 EPHA4 MAP1B RAC1 GPM6A THY1 NPTXR CLU SEPT7 APP PRNP GRIN1 DNER STMN4 SYT1 ATP2B1 HSP90AA1 GNAS HSP90AB1 CANX NDFIP1 SYT11 CRH [illegible] |
| 3.34448717619936E-13 | 39 | 1523 | Plasma membrane bounded cell projection part | GABRA1 PEBP1 SLC32A1 STMN2 HSPA8 GABRG2 EPHA4 MAP1B RAC1 GPM6A THY1 NPTXR CLU SEPT7 APP PRNP GRIN1 DNER STMN4 SYT1 ATP2B1 HSP90AA1 GNAS HSP90AB1 CANX NDFIP1 SYT11 CRH [illegible] |
| 3.6722896798768E-13 | 47 | 2214 | Plasma membrane bounded cell projection | STMN4 GABRA1 SYT1 PEBP1 SLC32A1 STMN2 HSPA8 GABRG2 EPHA4 STMN1 MAP1B SYT11 RAC1 GPM6A THY1 SLC6A1 SV2A NPTXR CD200 CLU SEPT7 APP PRNP GRIN1 DNER PKM ATP2B1 HSP90AA1 GNAS [illegible] |
| 1.00805392590675E-12 | 59 | 3479 | Extracellular space | MDH1 PKM ATP2B1 ATP6AP1 HSP90AA1 APLP2 DNAJA1 FTL GNAS PEBP1 HSP90AB1 PLD3 ALDOC CPE HSPA8 GAPDH LDHB STMN1 CLU SEPT7 CANX YWHAH ARF3 NARS ITM2B RAC1 TUBB2A APP ATP1B1 [illegible] |
| 1.00805392590675E-12 | 61 | 3693 | Extracellular region part | MDH1 PKM ATP2B1 ATP6AP1 HSP90AA1 APLP2 DNAJA1 FTL GNAS PEBP1 HSP90AB1 PLD3 ALDOC CPE HSPA8 GAPDH LDHB STMN1 CLU SEPT7 CANX YWHAH ARF3 NARS ITM2B RAC1 TUBB2A APP ATP1B1 [illegible] |
| 5.93605622231321E-12 | 40 | 1761 | Cytoplasmic vesicle part | SYT1 SCAMP1 SLC32A1 SV2A CLIP3 ITM2B RAC1 SERPINI1 GABRA1 SCG3 CPE TMEM30A GABRG2 EPHA4 NDFIP1 SYT11 CALM3 NSG2 CALM1 VAMP2 PKM ATP6AP1 HSP90AA1 APLP2 FTL HSP90AB1 PGRMC1 [illegible] |

**Table S2. Gene ontology analysis of the top-100 genes correlating with *ATP1A3* in the EN cluster in the infant cortex, Related to Figure 5**

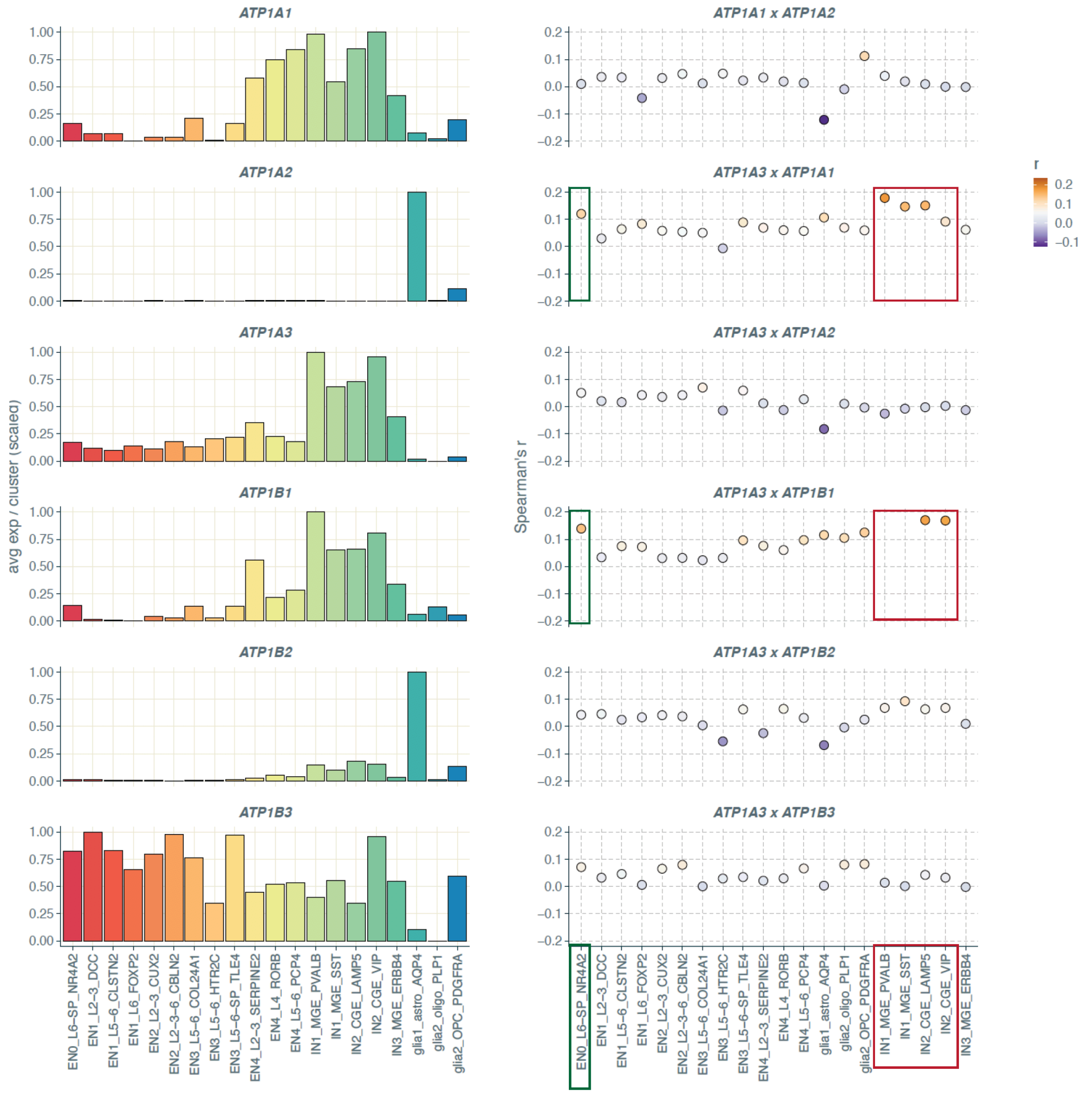


**Figure S5. Pair-wise correlation of *ATP1A1-3* in the infant human neocortex; Related to Figure 5**

**Figure S5. Pair-wise correlation of *ATP1A1-3* beta subunits in the infant human neocortex; Related to Figure 5**

*Left*, Bar graphs showing relative expression of *ATP1A1*, *ATP1A2* and *ATP1A3* (y-axis) across clusters. Gene expression (y-axis) was normalized by cell (divided by total UMIs/cell x 100k), then the mean value for each gene was aggregated by cluster and rescaled from 0 to 1. *Right,* Dot plots showing pairwise correlation coefficients between *ATP1A1, ATP1A2* and *ATP1A3* expression across clusters. Dots represent Spearmann's correlation coefficients (r), color coded by association strength. Of note, within ENs, the only cluster in which *ATP1A3* is coexpressed with *ATP1B1* significantly is layer 6 (SP_NR42A), labeled with green box. Also, note the positive correlation between expression of *ATP1A3* and *ATP1A1*, within the MGE derived cluster, most notably parvalbumin (PVALB), labeled with red box. EN, excitatory neuron; IN, inhibitory neuron.

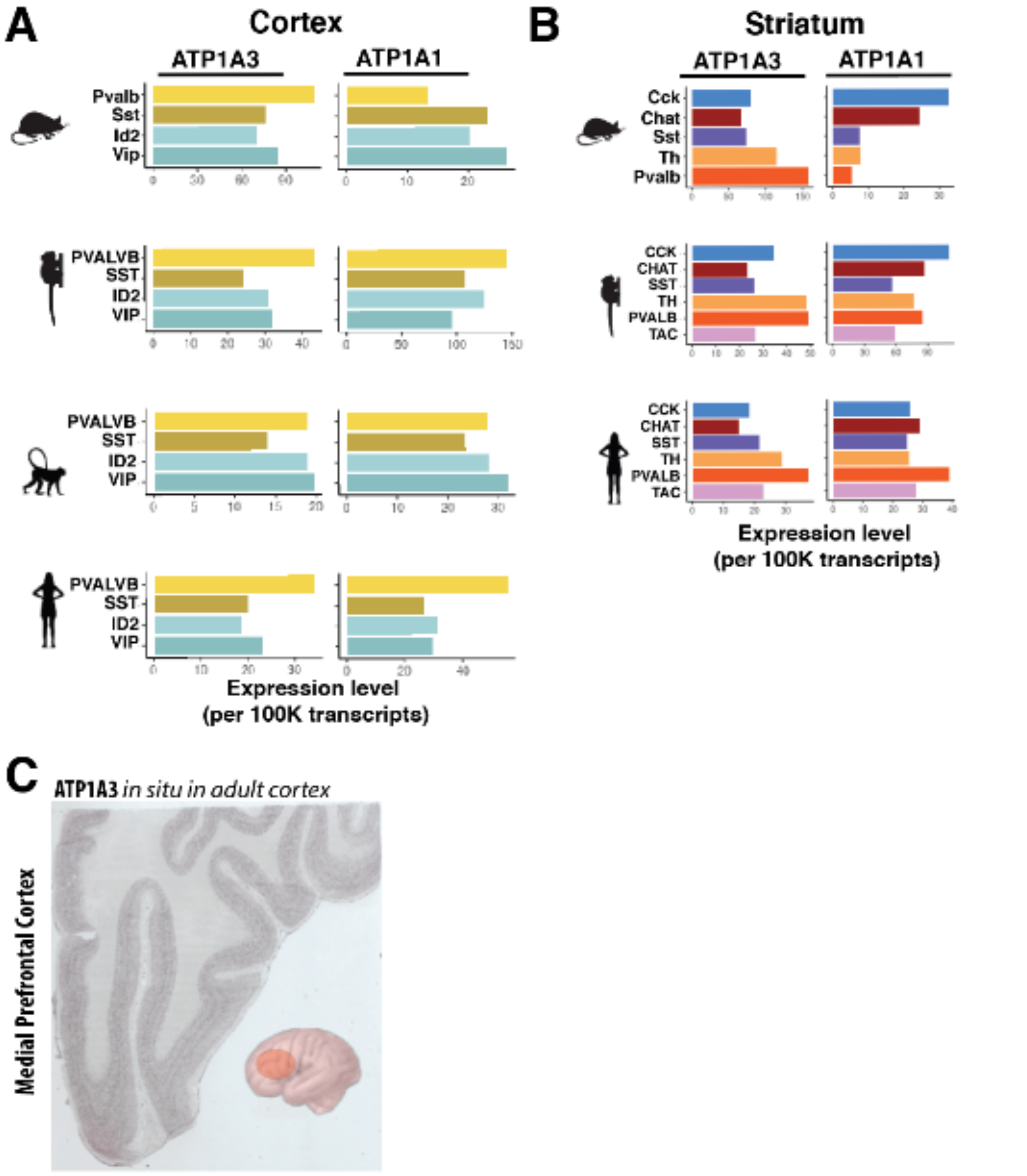


**Figure S6, Comparison of *ATP1A3* expression within cortical interneurons isolated from mice, macaque, and human; Related to Figure 5**

**Figure S6, Comparison of *ATP1A3* expression within cortical interneurons isolated from mice, macaque, and human; Related to Figure 5**

**(A)** Human vs. mouse, marmoset, and macaque comparing the four major neocortical interneuron classes: somatostatin ($SST^{+}$), parvalbumin ($PVALB^{+}$), Vasoactive intestinal peptide ($VIP^{+}$), or differentiation marker clade (ID2+). **(B)** Striatal interneurons colored by in one of the four major neocortical classes: cholecystokinin ($CCK^{+}$), choline acetyltransferase ($CHAT^{+}$), SST+, tyrosine hydroxylase ($TH^{+}$), $PVALB^{+}$, neuropeptide $TAC^{+}$. Scaled expression levels (number of transcripts per 100k) for *ATP1A3* and *ATP1A1*. Data from recently published collaborative study (Krienen et al, 2020). **(C)** *ATP1A3 in situ* demonstrates ubiquitous expression in the human adult cerebral cortex. (C) Image downloaded from the Allen Brain Atlas.

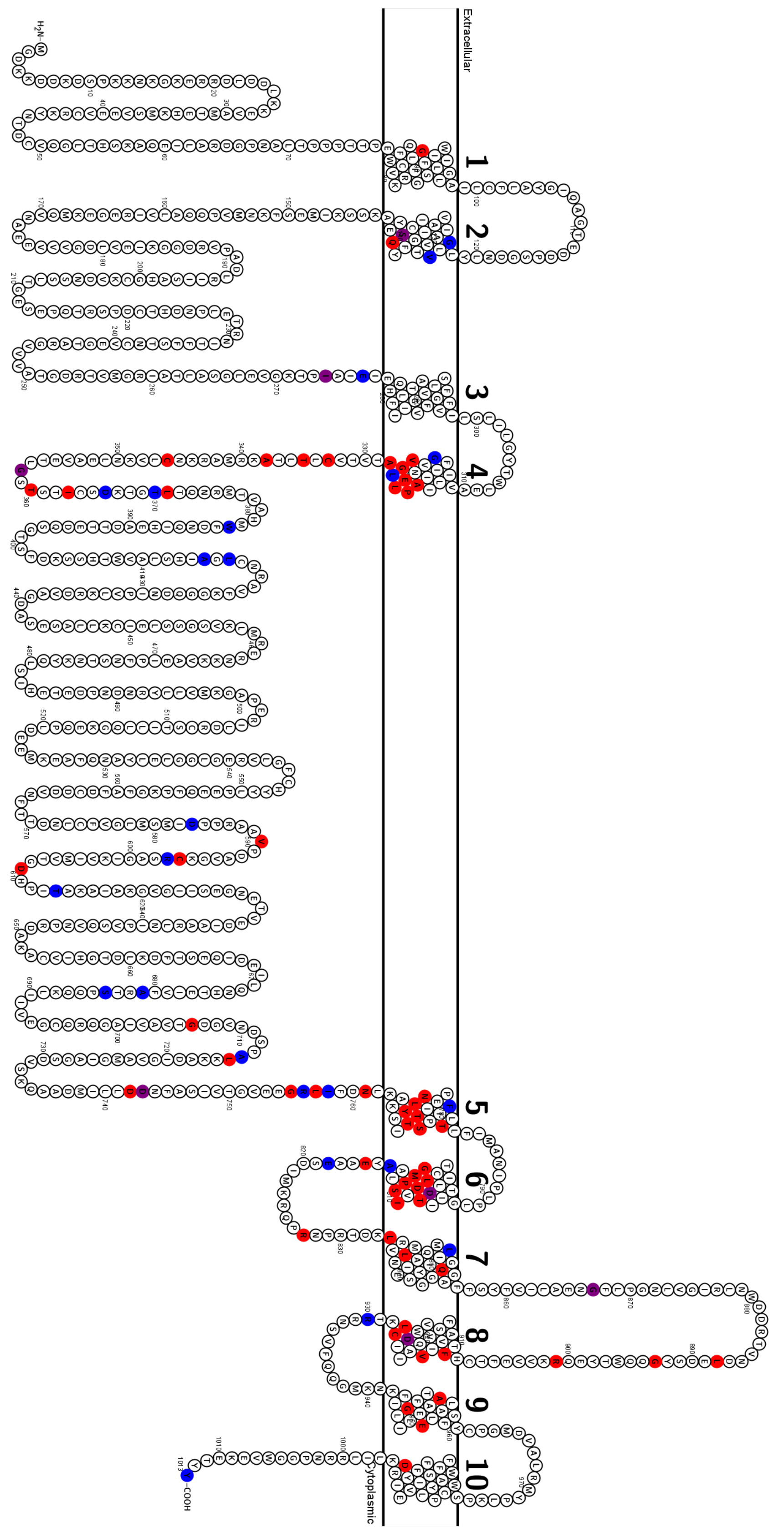


**Figure S7** Schematic of *ATP1A3* variants with respect to disease severity

**Figure S7** Schematic of *ATP1A3* variants with respect to disease severity.

**(A)** Phenotypes associated with *ATP1A3* are complex and overlapping, one way to organize them is by "severity" (i.e. age of onset), as done by Sweadner et al. 2019. Red corresponds to "severe", blue to "mild", and purple to "mixed" (e.g. different presentations with same mutation). Diseases like AHC and PMG have clear presentation in pre and early postnatal period, whereas patients of RDP may be asymptomatic until adulthood. Grouping mutations by the onset of associated symptoms in those patients suggests that mutations in the TM domains, and particularly in TMs 5-8 are more likely to lead to severe/early onset disease, and mutations in non-TM domains are more likely to lead to later onset *ATP1A3* related disease such as RDP. See Supplemental Table S3 for sources of each allele. See Figure 2C for disease specific mutations.

**Table S3: Summary of published *ATP1A3* variants associated with disease with original references and phenotypic features, See Figure 2C.**

| Sequence Variant (NM_152296.5) | Protein Variant | First Report | Severity | Documented Phenotype(s) |
|---|---|---|---|---|
| 55G>A | R19C* | Allocco 2019 | S | CH/Schizencephaly |
| 266delG | 89fs | Holze 2018 | S | HT/DD |
| 367G>C | G123R | Sampson 2016 | M | RDP |
| 385G>A | V129M | Smedemark-Margulies 2016 | M | COS/DD |
| 410C>A | S137Y | Heinzen 2012 | S | AHC |
| 410C>T | S137F | Heinzen 2012 | S | AHC |
| 409-411delGAG | S137del | Wilcox 2015 | M | RDP/PXD |
| 419A>T | Q140L | Heinzen 2012 | S | AHC |
| 420G>C or T | Q140H | Prange 2017 | S | RDP/EIEE |
| 821T>A | I274N | Rosewich 2012, Heinzen 2012 | S | AHC |
| 821T>C | I274T | de Carvalho Aguiar 2004 | M | RDP |
| 829G>A | E277K | de Carvalho Aguiar 2004 | M | AHC, RDP |
| 946G>A | G316S | Sweadner 2016 | M | RDP/CA |
| 958G>A | A320T | Trump 2016 | S | EIEE/PXD |
| 958G>C | A320P | ClinVar | | |
| 965T>A | V322D | Rosewich 2012 | S | AHC |
| 967C>T | P323S | Masoud 2017 | S | AHC |
| 968C>T | P323L | ClinVar** | | |
| 970G>C | E324Q | Panagiotakaki 2015, Viollet 2015 | S | AHC |
| 971A>G | E324G | Prange 2017 | S | AHC |
| 972G>C | E324D | Viollet 2015 | S | AHC |
| 974G>A | G325D | Lee 2014 | S | DD/CA |
| 977T>G | L326R | Panagiotakaki 2015, Viollet 2015 | S | AHC |
| 976-978delCTG | L327del | Kamm 2008 | M | RDP |
| 983C>A | A328D | Sweadner 2019 | S | |
| 998G>T | C333F | Heinzen 2012 | S | AHC |
| 1003A>C | T335P | Rosewich 2014a | S | AHC |
| 1004C>T | T335M | ClinVar | | |
| 1004C>A | T335K | ClinVar | | |
| 1012G>C | A338P | Gurrieri 2016 | S | AHC |
| 1036T>C | C346R | Marzin 2018 | S | HT/EIEE/DD |
| 1072G>T | G358C | Sasaki 2014 | S | EIEE/AHC |
| 1072G>A | G358S | Panagiotakaki 2015 | S | AHC |
| 1073G>T | G358V | Paciorkowski 2015 | S | EIEE |
| 1073G>A | G358D | Pereira 2015 | M | AHC/RDP |

| | | | | |
|---|---|---|---|---|
| 1079C>G | T360R | Prange 2017 | S | EIEE/CA/DYT |
| 1088T>A | I363N | Paciorkowski 2015 | S | EIEE/PXD/MC |
| 1088T>C | I363T | Kim 2020 | S | AHC/EIEE |
| 1096G>C | D366H | Sweadner 2019 | M | RDP |
| 1108G>A | T370A | Sweadner 2019 | M | |
| 1109C>A | T370N | Yang 2014, Rosewich 2014a | M | AHC, RDP |
| 1112T>C | L371P | Rosewich 2012 | S | AHC |
| 1123C>T | R375C | ClinVar | | |
| 1124G>A | R375H | ClinVar | | |
| 1144T>C | W382R | Rosewich 2012 | M | AHC/RDP |
| 1244C>A | A415D | Sweadner 2019 | M | |
| 1250T>C | L417P | Rosewich 2014a | M | RDP |
| 1387G>A | R463C* | Allocco 2019; Aryastarkhova 2019 | S | CH/Schizencephaly, RDP** |
| 1747G>T | D583Y | Nicita 2016(Nicita et al., 2016)35 | M | PXD/RDP |
| 1765G>T | V589F | Richards 2018; Tran 2020 | S | EIEE/AHC/RDP |
| 1786T>C | C596R | Viollet 2015 | S | AHC |
| 1790G>C | R597P | Wenzel 2017 | M | RDP |
| 1795G>A | A599T | ClinVar | | |
| 1825G>T | D609Y | Marzin 2018 | S | HT/DD/EIEE |
| 1838C>T | T613M | de Carvalho Aguiar 2004; Balint 2019 | M | RDP, PXD |
| 2041G>A | A681T | Torres 2018 | M | DD/ASD[a] |
| 2051C>T | S684F | Svetel 2010 | M | RDP |
| 2116G>A | G706R | Yang 2014 | S | AHC |
| 2140G>A | A714T | Meijer 2016 | M | RDP |
| 2144T>C | L715P | Panagiotakaki 2015 | S | AHC |
| 2224G>T | D742Y | Marzin 2018 | S | EIEE/PXD |
| 2227G>C | D743H | Meijer 2016 | M | RDP |
| 2266-2268delGAC | D743del | Schirzini 2018 | S | EIEE/AHC/HT |
| 2263G>A | G755S | Heinzen 2012 | S | AHC |
| 2263G>T | G755C | Rosewich 2012 | S | AHC |
| 2264G>T | G755V | Viollet 2015 | S | AHC |
| 2264G>C | G755A | Sasaki 2014 | S | AHC |
| 2266C>T | R756C | Dard 2015 | M | RECA |
| 2267G>A | R756H | Brashear 2012 | M | RECA |
| 2267G>T | R756L | Yano 2017 | M | RECA |
| 2270T>C | L757P | Rosewich 2014a | S | AHC |
| 2272A>T | I758F | Sweadner 2019 | M | |
| 2273T>G | I758S | de Carvalho Aguiar 2004 | M | RDP |
| 2281A>C | N761H | Viollet 2015 | S | AHC |
| 2302T>C | Y768H | Viollet 2015 | S | AHC |

| | | | | |
|---|---|---|---|---|
| 2303T>C | Y768C | Viollet 2015 | S | AHC |
| 2305A>C | T769P | Viollet 2015 | S | AHC |
| 2309T>G | L770R | Yang 2014 | S | AHC |
| 2312C>A | T771N | Sasaki 2014 | S | AHC |
| 2312C>T | T771I | Yang 2014 | S | AHC |
| 2314A>C | S772R | Panagiotakaki 2015, Viollet 2015 | S | AHC |
| 2316C>A | S772R | Rosewich 2012 | S | AHC |
| 2316C>G | S772R | Yang 2014 | S | AHC |
| 2317A>C | N773H | Viollet 2015 | S | AHC |
| 2318A>G | N773S | Heinzen 2012 | S | AHC |
| 2318A>T | N773I | Rosewich 2012 | S | AHC |
| 2318A>C | N773T | Yang 2015 | S | AHC |
| 2323C>A | P775L | ClinVar (2) | | |
| 2324C>T | P775T | ClinVar | | |
| 2332A>C | T778P | Gasser 2020 | S | RDP/EP |
| 2338T>C | F780L | de Carvalho Aguiar 2004 | M | RDP |
| 2401G>A | D801N | Heinzen 2012 (>300) | S | AHC, AHC/RDP |
| 2401G>T | D801Y | de Carvalho Aguiar 2004 | M-S | AHC, RDP |
| 2401G>C | D801H | ClinVar | | |
| 2402A>T | D801V | Panagiotakaki 2015 | S | AHC |
| 2403T>A | D801E | Hoei-Hansen 2014 | S | AHC |
| 2405T>G | L802R | Zúñiga-Ramírez 2019 | S | PXD |
| 2405T>C | L802P | Yang 2014 | S | AHC |
| 2408G>A | G803D | Gall 2017 | S | CA/DYT/DD/MC/HT |
| 2411C>T | T804I | Ulate-Campos 2014, Rosewich 2014b | S | AHC |
| 2413G>A | D805N | Viollet 2015 | S | AHC |
| 2413G>C | D805H | Yang 2014 | S | AHC |
| 2415C>G | D805E | Rosewich 2014 | S | AHC |
| 2417T>G | M806R | Heinzen 2012 | S | AHC |
| 2417T>A | M806K | Yang 2014 | S | AHC |
| 2423C>T | P808L | Yang 2014 | S | AHC |
| 2428A>T | I810F | Rosewich 2014 | S | AHC/RDP |
| 2429T>G | I810S | Heinzen 2012 | S | AHC |
| 2429T>A | I810N | Yang 2014 | S | AHC |
| 2431T>C | S811P | Heinzen 2012 | S | AHC |
| 2438C>T | A813V | Kubota 2017 | M | COS/ASD |
| 2443G>A | E815K | Heinzen 2012 ( >200) | S | AHC, AHC/COS |
| 2452G>A | E818K | Demos 2014 | M | CAPOS, AHC/CAPOS |
| 2479A>T | R827W | Gasser 2020 | S | AHC |
| 2501T>C | L834S | Yang 2015 | S | AHC |

| | | | | |
|---|---|---|---|---|
| 2516T>C | L839P | Yang 2014 | S | AHC |
| 2542+1G>A | SPLICE | Viollet 2015 | S | AHC |
| 2542+2T>C | SPLICE | Viollet 2015 | S | AHC |
| 2552A>G | Q851R | Masoud 2017, This study | S | AHC, PMG |
| 2552A>C | Q851P | Yang 2015 | S | AHC/EIEE |
| 2558T>G | L853R | Rodriguez-Quiroga 2016 | M | AHC, RDP, AHC/RDP |
| 2600G>A | G867D | Rosewich 2014b | M | AHC/RDP |
| 2600G>A | G867N | Sweney 2015*** | S | AHC, RDP |
| 2663T>C | L888P | Panagiotakaki 2015, This study | S | AHC, PMG |
| 2677G>A | G893R | Yang 2014 | S | AHC |
| 2702G>C | R901T | Viollet 2015 | S | AHC |
| 2702G>T | R901M | This study | S | PMG |
| 2736-2738delCTT | F913del | Ishihara 2019 | S | AHC/HT/EIEE |
| 2755- 2757delGTC | V919del | Heinzen 2012 | S | AHC |
| 2767G>T | D923Y | Rosewich 2012 | S | AHC |
| 2767G>A | D923N | Zanotti 2008 | M | AHC/RDP |
| 2767G>T | D923T | Sweney 2015*** | S | AHC |
| 2810T>C | L924P | Aryastarkhova 2019, This study | S | EIEE/MC, PMG |
| 2780G>A | C927Y | Ishii 2013 | S | AHC |
| 2780G>T | C927F | Sasaki 2014 | S | AHC |
| 2781C>G | C927W | Ulate-Campos 2014 | S | AHC |
| 2788C>T | R930W | Meijer 2016 | M | RDP |
| 2839G>A | G947R | Heinzen 2012 (>30) | S | AHC, EIEE, HT/AHC |
| 2839G>C | G947R | Heinzen 2012 (>50) | S | AHC, EIEE, HT/AHC |
| 2851G>A | E951K | Panagiotakaki 2015, Viollet 2015 | S | AHC, AHC/RDP |
| 2864C>A | A955D | Heinzen 2012 | S | AHC |
| 2960+1G>A | SPLICE | This study | S | PMG |
| 2974G>C | D992H | Sweadner 2019 | S | AHC |
| 2974G>T | D992Y | Heinzen 2012 | S | AHC |
| 3191-3193dupTAC | Y1013YY | Blanco-Arias 2009 | M | RDP |

*Seen in recessive case
**Atypical presentation/responsive to L-DOPA treatment
***Could not locate original report, earliest found reference

**Methods**: Based on methods and criteria used by Sweadner et al. 2019[17]; see supplementary table e-4. Severity was defined by age of onset of symptoms (S = severe/neonatal onset, M = milder/later onset). Phenotypes were annotated based on review of published clinical information in the original report of each allele, classified with descriptions/keywords. Some mutations were originally reported on a different transcript, but were adjusted to reflect NM_152296.5, the canonical *ATP1A3* transcript, as done by Sweadner et al. 2019. Annotations on Protter (http://wlab.ethz.ch/protter/) on Figure 2C and Figure S6.

**Phenotype**

| | |
|---|---|
| **HT** | Hypotonia (diffuse) |
| **MC** | Microcephaly |
| **DD** | Developmental Delay |
| **EIEE** | Early Infantile Epileptic Encephalopathy |
| **AHC** | Alternating Hemiplegia of Childhood |
| **RDP** | Rapid Onset Dystonia-Parkinsonism |
| **RECA** | Recurrent Episodes of Cerebellar Ataxia (Fever Induced) |
| **CAPOS** | Cerebellar Ataxia, Areflexia, Pes Cavus, Optic Atrophy, and Sensorineural Hearing Loss |
| **PXD** | Paroxysmal Dystonia |
| **CH** | Congenital Hydrocephalus |
| **PMG** | Polymicrogryria |
| **COS** | Childhood Onset Schizophrenia |
| **CA** | Cerebellar Ataxia (or confirmed cerebellar atrophy on MRI) |
| **DYT** | Dystonia/Generalized |
